# Structural basis for monobody OP-4 binding to open-form adenylate kinase

**DOI:** 10.64898/2026.08.04.742914

**Authors:** Naoki Orito, Ibuki Nakamura, Katsuhisa Yoshihara, Kosuke Onishi, Sachiko Toma-Fukai, Hiroaki Matsuura, Shun-ichi Tanaka, Takashi Matsuo

## Abstract

Monobodies are fibronectin type-III-based binding proteins that specifically bind target proteins and regulate their functions. We previously identified monobodies that selectively recognize either the OPEN or CLOSED conformation of adenylate kinase (Adk), revealing that monobodies can discriminate distinct conformational states of a target protein. However, the molecular basis of OPEN-form recognition has remained unclear because the structure of the complex between an OPEN-form-specific monobody and Adk had not been determined. To address this issue, we determined the crystal structure of the complex between Adk and the OPEN-form-specific monobody OP-4, employing hierarchical clustering analysis of X-ray diffraction datasets. The structure revealed that OP-4 binds to the surface formed by the expanded LID and CORE domains of Adk. Mutations in the interface residues reduced the OP- 4-binding affinity, indicating that the crystallographically identified interface is also relevant in solution. In particular, R123 mutations markedly impaired OP-4 binding. Molecular dynamics simulations further suggested that the R123–D159–R156 hydrogen-bond network is retained in solution and may contribute to efficient complex formation.

These findings establish the structural basis for monobody OP-4 binding to open-form Adk and identify the R123-centered interaction network as a key determinant of complex formation.

**Graphical Abstract:** 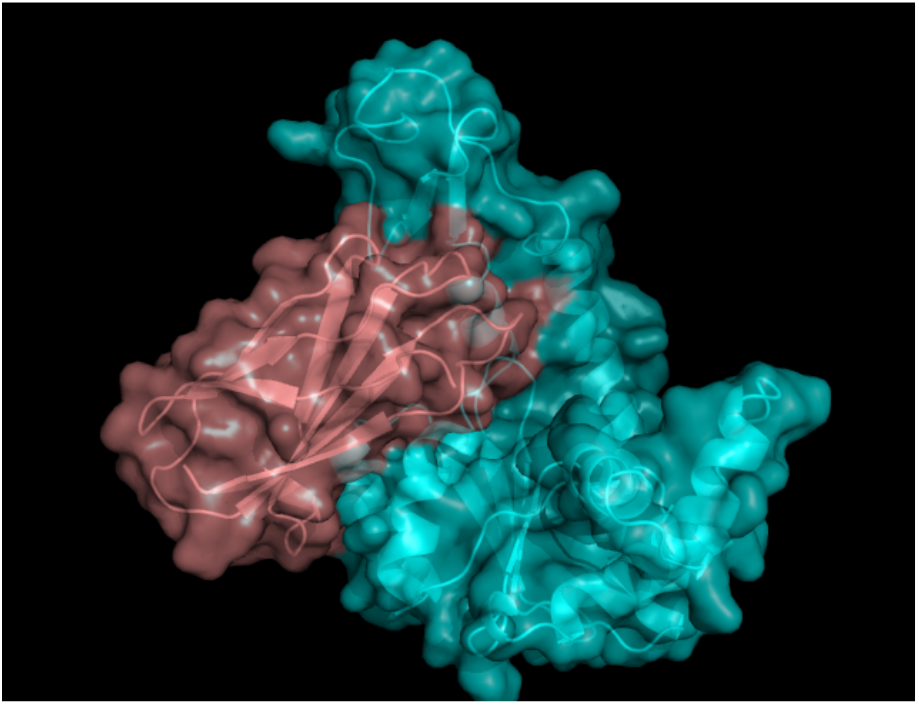

**Highlights:**

- The crystal structure of the OPEN-form adenylate kinase/monobody OP-4 complex was successfully determined.
- The crystal structure revealed that OP-4 binds to surface formed by the expanded LID and CORE domains in adenylate kinase.
- Isothermal titration calorimetry measurements for adenylate kinase mutants confirmed that the binding modes observed in solution are consistent with those observed in the crystal structure.
- Molecular dynamics simulations suggested that the R123–D159–R156 hydrogen bond network prior to OP-4 binding is important for complex formation.

## Introduction

Monobodies are synthetic binding proteins with a molecular scaffold of the 10th human fibronectin type-III (FN3) domain (*ca*. 10 kDa), functioning as an antibody mimetic that binds specifically to target proteins (Fig. 1(a)) [1–3]. Because of their potential as alternatives to conventional antibodies, monobodies have been developed against a variety of disease- associated proteins [4–9].

**Fig. 1.**
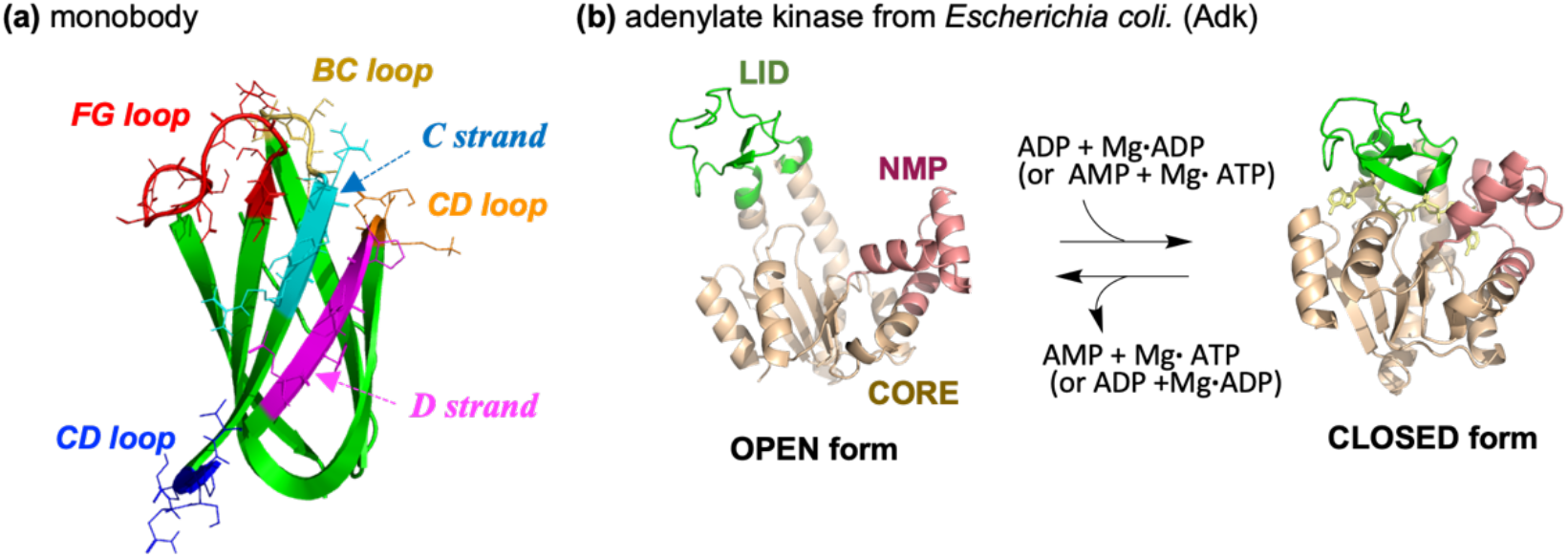
Structures of FN3-based monobody and adenylate kinase from *Escherichia coli*. (Adk); (a) Structure of FN3-based monobody (PDB: 1FNA); (b) OPEN- and CLOSED-form structures of Adk (PDB: 4AKE (OPEN-form), 1AKE (CLOSED- form)); In figure (a), diversified regions in “side-and-loop” library are indicated. In figure (b), the CORE domain (residues 1–29, 68–117, 161–214), NMP domain (residues 30–67), and LID domain (residues 118–160) are marked in wheat, light red, and green, respectively.

Diverse monobodies targeting various proteins can be created through random mutations on the protein surface using phage or yeast-surface display systems [3, 10]. Two types of monobody libraries are known for monobody diversification [2]. One is a “loop-only library,” which involves random mutations at loop regions (*BC*-, *DE*-, and *FG*-loops). The other is a “side-and-loop library,” which is constructed by mutations at strand regions (*C*- and/or *D*-strands) as well as the loop region. Structural studies have shown that the mutation patterns in these libraries determine the binding modes of the resulting monobodies [4, 5, 11, 12].

Complex formation of monobodies with their target proteins is achieved through a wide range of protein-protein interactions. Such interactions are often favored when the binding interface is pre-organized, minimizing conformational rearrangements and the associated entropic penalties [13]. In this context, we became interested in the binding mechanism of monobodies toward proteins that have conformational fluctuations in solution; thus, we have investigated the binding mechanism of monobodies for adenylate kinase from *E. coli* (Adk; Fig. 1(b); see the amino acid sequence in Table S1), an enzyme that mediates phosphoryl transfer between nucleotides (Mg•ADP + ADP ⇔ Mg•ATP + AMP) [14, 15].

Adk undergoes an OPEN/CLOSED conformational transition during nucleotide binding and release, where NMP and LID domains (see Fig. 1(b)) repeatedly approach and dissociate owing to a hinge motion at CORE domain [16]. Additionally, Adk is known to transiently adopt CLOSED-form-like conformations in solution without nucleotides, owing to its backbone flexibility (denoted as “conformation population-shift”) [17, 18].

Previously, we obtained four monobodies from a “side-and-loop library”: one monobody (CL-1) selectively recognizes the CLOSED-form Adk, whereas the other three clones (OP-2, -3, and -4) bind to the OPEN-form [14]. Among the four monobodies, OP-4 is particularly intriguing because OP-4 exhibits the highest binding affinity and inhibits kinase activity (see amino acid sequence of OP-4 in Table S1). Our previous report, including size exclusion column chromatography/small-angle X-ray scattering (SEC-SAXS) analysis, suggested that OP-4 binds to Adk while retaining an overall OPEN-form conformation [14]. The study using ^31^P-NMR spectroscopy also supported this idea: ATP, a nucleotide substrate for Adk, can bind to Adk even in the OP-4-bound form, suggesting the maintenance of an OPEN-form-like structural state [15]. However, crystallographic analysis of the Adk:OP-4 complex has been hampered by heterogeneity in the diffraction data, preventing us from obtaining a high-resolution structure. Thus, the residue-level interaction mechanism for the OP-4 binding to Adk has been remained unclear.

Accordingly, we performed hierarchical clustering analysis (HCA)[19] in this study to resolve the crystal structure of the Adk:OP-4 complex. HCA is useful for classifying heterogeneous diffraction datasets into several distinct clusters with various degrees of isomorphism. As a result, we successfully determined the crystal structure of OP-4-bound Adk. In this paper, we report the crystal structure of the Adk:OP-4 complex determined via HCA-assisted analysis. Furthermore, we performed mutational, thermodynamic, and molecular dynamics (MD) analyses to evaluate whether the interactions observed in the crystal structure are functionally relevant in solution. Combined with crystallographic and solution-state analyses, the structural information provides residue-level insights into the molecular mechanism underlying Adk recognition by OP-4.

## Materials and Methods

### Materials

All chemicals and column supports were purchased from commercial sources and used as received. wild-type Adk (wtAdk) was expressed in *E. coli.* HB101 with wtAdk- harboring pEAK91 plasmid and purified by the previously reported method [20, 21]. Monobody OP-4 was expressed in *E. coli.* BL21(DE3) with OP-4-harboring pHFT2 plasmid and firstly prepared as a protein with a histidine-tag at *N*-terminus as previously reported [3, 22]. The monobody used in this study was obtained by histidine-tag cleavage by TEV protease according to previous reports [3, 20, 22, 23]. Adk mutants were obtained as described below. Crystals of the OPEN-form Adk:OP-4 complex were prepared as described below.

### Instruments

Protein purification and chromatographic analyses were performed using an ÄKTA FPLC or ÄKTAprime chromatography system (GE healthcare). Isotherm titration calorimetric (ITC) measurements were performed using a Malvern MicroCal^TM^ PEAQ-ITC or ITC_200_ microcalorimeter. Circular dichroism (CD) spectra were collected using a JEOL J-820 circular dichroism spectropolarimeter. X-ray diffraction data were collected at the BL45XU beamline of SPring-8, Japan.

### Preparation and purification of Adk mutants

pEAK91-based plasmids coding Adk mutants were prepared by polymerase chain reactions for the parent plasmid coding wtAdk using TOYOBO KOD-plus mutagenesis kit. These plasmids were used to transform *E. coli.* HB101. Adk mutants except for R123D and R123E were obtained in the same manner as wtAdk[20, 21]. Briefly, extracts from *E. coli.* bodies were dissolved in 50 mM Tris-HCl (pH 7.5 at 25°C) containing 1 mM 2-ME and passed through a DEAE column. The flow path fractions were loaded onto a Blue Sepharose (Cytiva). The fraction components eluted with a linear gradient (A: 50 mM Tris-HCl with 1 mM 2-ME (pH 7.5); B: 50 mM Tris-HCl with 1 mM 2-ME and 1 M KCl (pH 7.5)). After concentration, the protein solution was loaded on a HiLoad Sephadex G75 with elution of 50 mM Tris-HCl with 1 mM TCEP (pH 7.5 at 4°C) to obtain purified mutant proteins. For the purification of R123D and R123E mutants, a HiTrap Q column (5 mL; Cytiva) was used in place of a Blue Sepharose column. The purities of prepared mutants were checked by SDS-PAGE. All Adk mutants can be stocked in 20 mM Tris-HCl (pH 8) that contains 1 mM TCEP and glycerol (40%(v/v) in final) at –20°C for at least 6 months. Immediately before use, these additives were removed by gel filtration using HiTrp desalting column (Cytiva).

### Crystallization of the OPEN-form Adk:OP-4 complex

To a concentrated solution of wtAdk (500 μM) and OP-4 (550 μM) in 20 mM Tris-HCl buffer (pH 8.0)). Crystals of the OPEN-form Adk:OP-4 complex for X-ray diffraction were prepared by hanging-drop vapor-diffusion method. The crystals were grown in 2 μL solutions that contained 1 μL of the complex solution and 1 μL of reservoir solution (0.1 M Tris-HCl pH 8.5 with 0.2 M Li_2_SO_4_, 30%(w/v) polyethylene glycol 4000, and 20%(v/v) glycerol), equilibrated against 100 μL of the reservoir solution at 20°C over 8 days. Crystals were secured on cryo-loops and flash-frozen at 100K using a nitrogen cryosystem.

### X-ray diffraction data collection and analyses

X-ray diffraction data were collected using the ZOO automated data collection system [24]. Each diffraction dataset was collected over a total oscillation range of 360° using the helical scanning method. Images were split into 5° sub-datasets to separate heterogeneous structures in the crystals in combination with the following HCA analysis. All diffraction datasets were automatically processed by the KAMO automated data reduction pipeline [25].

### Hierarchical clustering analysis (HCA) of X-ray diffraction data

HCA was performed as described previously [19] to classify diffraction datasets based on their isomorphism. The classified datasets were visualized using a dendrogram, where the vertical distance (Ward linkage distance) reflects the degree of dissimilarity; thus, datasets connected at smaller distances are considered more isomorphous.

### Crystallographic analysis of the OPEN-form Adk:OP-4 complex

Several datasets (clusters 65, 72, and 77–82) were selected for structural determination based on molecular replacement analyses, using PHENIX program [26]. Among the classified datasets (*i.e.*, clusters 1–81), the dataset with the highest completeness (cluster 81) was subjected to model refinement using phenix.refine in PHENIX [26], with manual inspection and modification in conjunction with the CCP4 program COOT [27]. The *φ*–*ϕ* angles of >90% of the residues in the structure of the Adk:OP-4 complex are in the most favored regions of the Ramachandran plot as assessed by MolProbity [28]. Phasing and refinement statistics are listed in Table S2. The final atomic coordinates and structure-factor amplitudes of the Adk:OP-4 complex were deposited in the Worldwide Protein Data Bank (wwPDB; http://www.wwpdb.org; PDB: 44AY) and the Protein Data Bank Japan at the Institute for Protein Research, Osaka University (Suita, Osaka, Japan; PDBj; http://www.pdbj.org/).

### CD spectral measurements of Adk mutants

Circular dichroism (CD) spectra of Adk mutants were measured for a Adk mutant (5 μM) in 5 mM potassium phosphate buffer (pH 8.0) at 25°C.

### Isothermal titration calorimetry (ITC) measurements for complexation of wtAdk and Adk mutants with monobodies

Isothermal titration calorimetry (ITC) measurements were performed at 25°C. A solution of an Adk mutant in 20 mM Tris-HCl buffer (pH 8.0) containing 150 mM NaCl was incubated in the sample cell (200 μL) and titrated with OP-4 (40 μL) with each injection of 2 μL with a 4 s duration, and a 150 s interval. The concentrations of an Adk mutant in the sample cell were set to 10 or 30 μM (30 μM for R123K, F137A, and D159K mutants; 10 μM for wtAdk and other mutants). The concentrations of OP-4 in the titration syringe were set to 110 or 730 μM (730 μM for R123K, F137A, and D159K mutants; 110 μM for wtAdk and other mutants). The obtained raw data were corrected by ligand heats of dilution and integrated using MicroCal PWAQ-ITC analysis software. The heat data were analyzed using a single-site binding model.

### Size exclusion chromatography (SEC) measurements

For R123X Adk mutants (X = A, D, or E), a solution of an Adk mutant with or without OP-4 (15 μL; [Adk] = 30 μM, [OP-4] = 0 or 42 μM) was prepared in 20 mM Tris-HCl (pH 8.0) containing 150 mM NaCl and loaded onto a Supderdex 75 5/150 GL column (Cytiva). The sample was eluted at a flow rate of 0.35 mL/min and detected at 280 nm.

### Molecular dynamic (MD) simulations

YASARA structure molecular modeling software package (Ver.25.1.13) [29] was used for MD simulations. MD calculations were performed using AMBER14 force field, an aqueous solution model with 0.9% NaCl*aq* ion concentration, point charges assigned at pH = 8.0 at 298.15 K, and additional Na^+^ or Cl^−^ for charge neutralization in a cubic cell boundary defined at 10 Å from the protein surface. The simulations were continued for 106 ns, where the change in Cα-RMSDs (root-mean-square deviations) reached a deviation within ± 1 Å. The X-ray crystallographic structure (PDB: 4AKE) was used as the initial structure of wtAdk. For Adk mutants, the initial structures were constructed by computational mutation of the wtAdk structure using YASARA. The trajectory was sampled every 100 ps. For each frame, inter-sidechain distances (ISDs), distances between predefined sidechain atom groups, were calculated. For residue pairs, atoms used for calculations are the guanidinium nitrogen atoms (NE, NH1, and NH2) in an arginine residue, the carboxylate oxygen atoms (OD1 and OD2) in an aspartic acid residue, the amine nitrogen atom (NZ) in a lysine residue, and the methyl carbon in alanine. Occupancy (OCC) values were obtained by counting the number of frames that satisfied the distance criterion (≤3.5 Å) and dividing by the total number of analyzed frames.

### Comparison with SAXS data

The previously acquired SAXS data were used for the calculation of electron density map. The electron density map was constructed on DENSSWeb (https://denss.ccr.buffalo.edu/submit) [30] in the same manner as our previous study [14]. Electron density map figures were created using ChimeraX (ver. 1.10.1) [31]. Structural fitting of the electron density map determined by DENSS and the crystal structure was performed using AutoFit in ChimeraX.

## Results

### HCA of polymorphic diffraction data for the Adk:OP-4 complex

After processing the diffraction data and grouping the equivalent cell group using KAMO system to remove data with low completeness, 83 of 5° sub-datasets were classified using HCA to sort various degrees of isomorphism. The resulting dendrogram is shown in Fig. 2. Among the clusters analyzed (clusters 65, 72, and 77–82), cluster 81 yielded the highest completeness, removing a secondary large data group (cluster 80). Accordingly, the dataset from cluster 81 was used for subsequent model refinement.

**Fig. 2.**
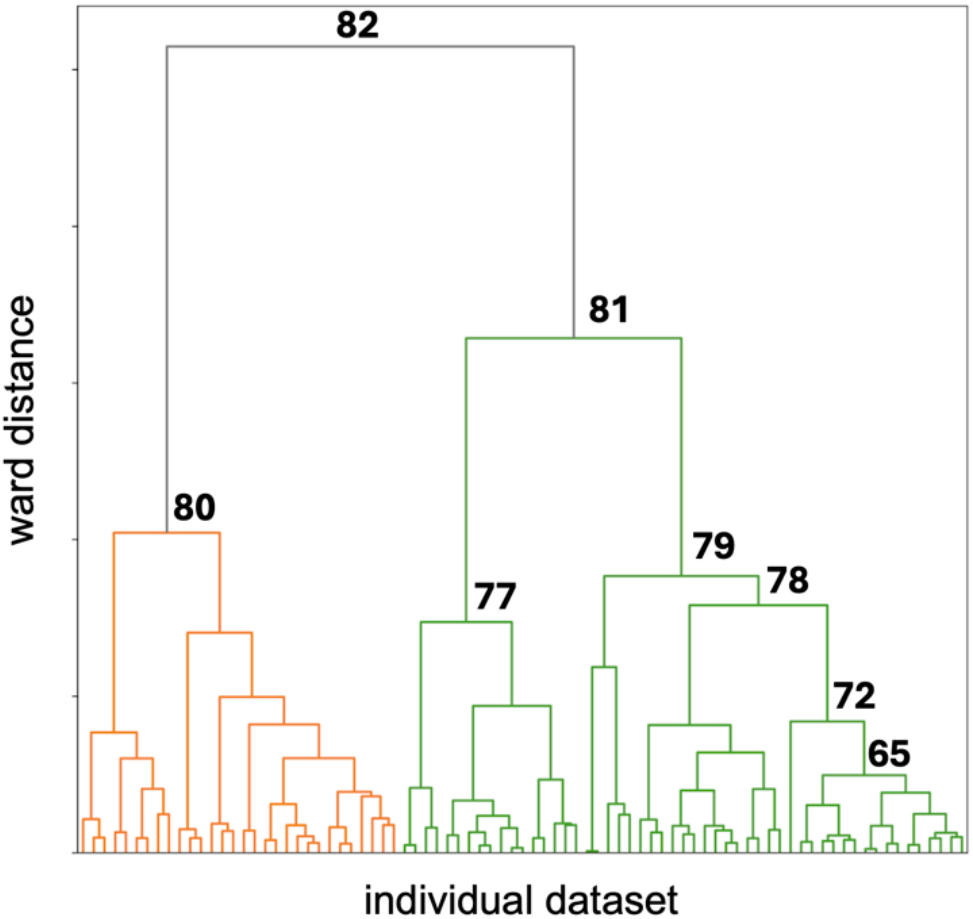
Dendrogram obtained by hierarchical clustering analysis (HCA) of polymorphic diffraction datasets for the OPEN-form Adk:OP-4 complex. The vertical distance represents the degree of dissimilarity between datasets.

### Crystal structure of the OPEN-form Adk:OP-4 complex determined from cluster 81 dataset

Fig. 3 shows the crystal structure of the OPEN-form Adk:OP-4 complex determined from cluster 81 dataset with a resolution of 2.53 Å. The structure indicated the OP-4 binding to the surface formed by the expanded LID and CORE domains, where *FG*-loop and *C*-strand in OP-4 are associated with the complex formation. In addition, *D*-strand in OP-4 also participates in the binding to Adk with “side” binding fashion. The OP-4 binding site shown in the crystallographic structure differs from that in the binding model proposed in our previous SEC-SAXS study.

**Fig. 3.**
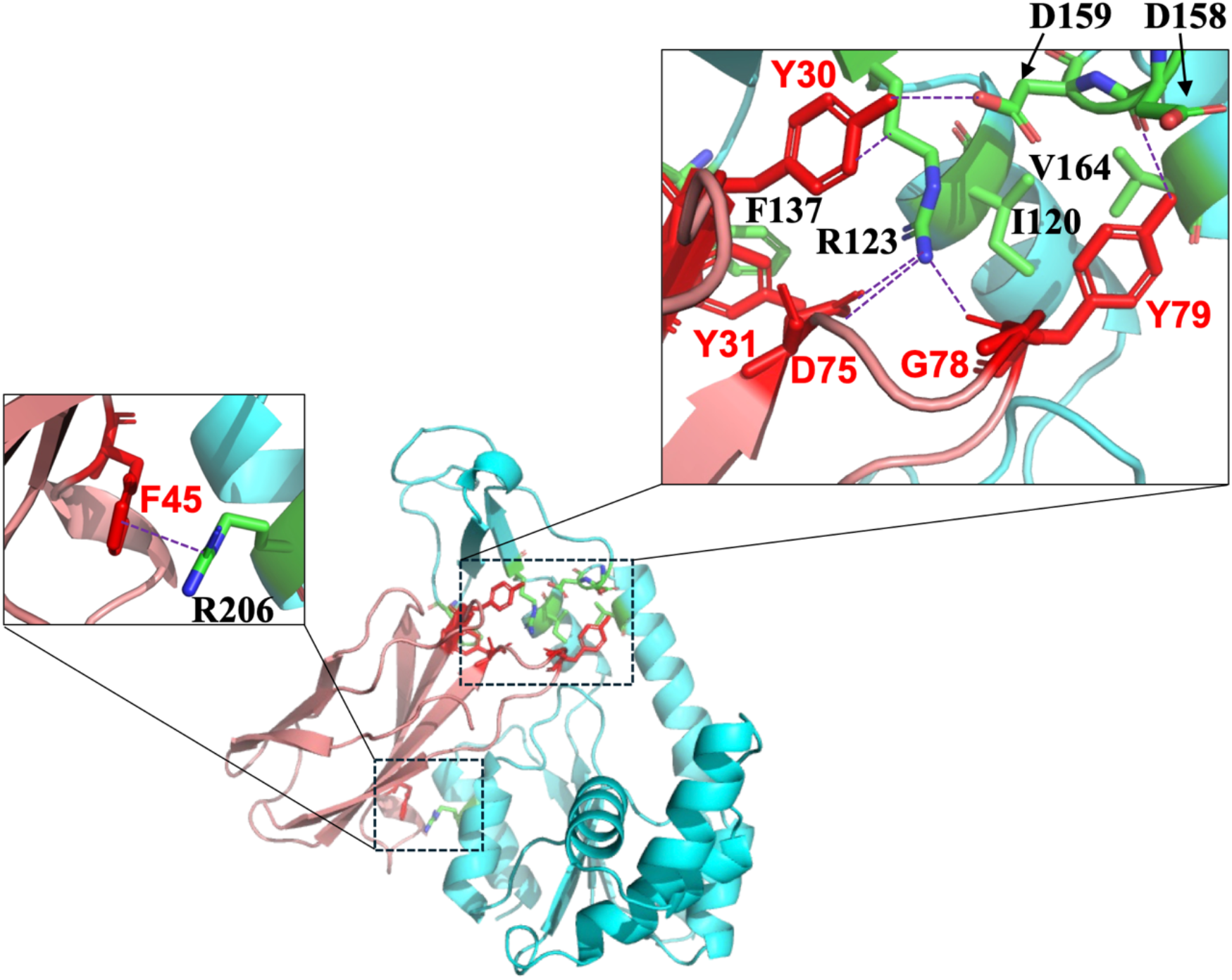
X-ray crystallographic structure of the OP-4:Adk complex (PDB: 44AY). The backbones of OP-4 and Adk are depicted in light red and light cyan color, respectively. Amino acid residues associated with complex formation are marked in sticks. The structural analysis was performed using the dataset cluster 81.

Table 1 lists amino acid residues involved in the complex formation. R123 and F137 on helix 6 (α6) in Adk interacts with Y30, Y31, D75, and G78 in OP-4. Especially, R123 has several interaction modes with OP-4. The distance between F137 in Adk and Y31 in OP-4 is identified as the existence of π– π stacking between them. D159 on β-strand 6 (β6) in Adk participates in the complex formation through the interaction with Y30 in OP-4. A hydrophobic core is formed between I120/V164 in Adk and Y79 in OP-4 through sidechain contacts with distances of < 5 Å. A cation–π interaction occurs between R206 in Adk and F45 on CD-loop of OP-4.

**Table 1.**
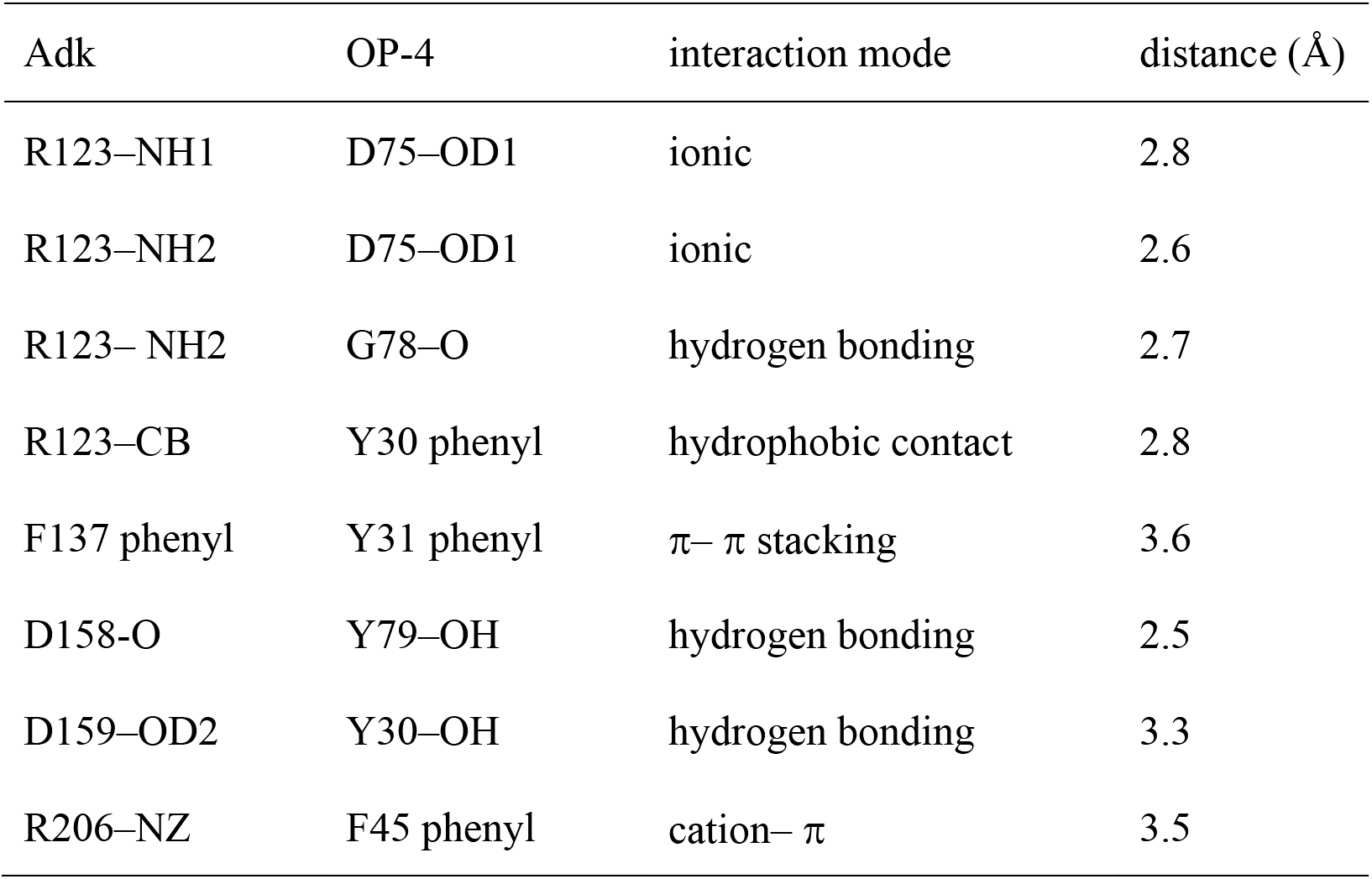
Interactions between wtAdk and OP-4 in their complex.

| Adk | OP-4 | interaction mode | distance (Å) |
| --- | --- | --- | --- |
| R123–NH1 | D75–OD1 | ionic | 2.8 |
| R123–NH2 | D75–OD1 | ionic | 2.6 |
| R123–NH2 | G78–O | hydrogen bonding | 2.7 |
| R123–CB | Y30 phenyl | hydrophobic contact | 2.8 |
| F137 phenyl | Y31 phenyl | π–π stacking | 3.6 |
| D158–O | Y79–OH | hydrogen bonding | 2.5 |
| D159–OD2 | Y30–OH | hydrogen bonding | 3.3 |
| R206–NZ | F45 phenyl | cation–π | 3.5 |

### CD spectra of Adk mutants

The Adk mutants prepared for elucidating the interaction modes in solution are listed in Table 2. As mutation sites, residues R123, F137, D158, and D159 were selected, which face to OP-4 at the boundary of LID and CORE domains, The CD spectra of these mutants are shown in Fig. 4. Spectral shapes of these mutants were similar to that of wtAdk.

**Fig. 4.**
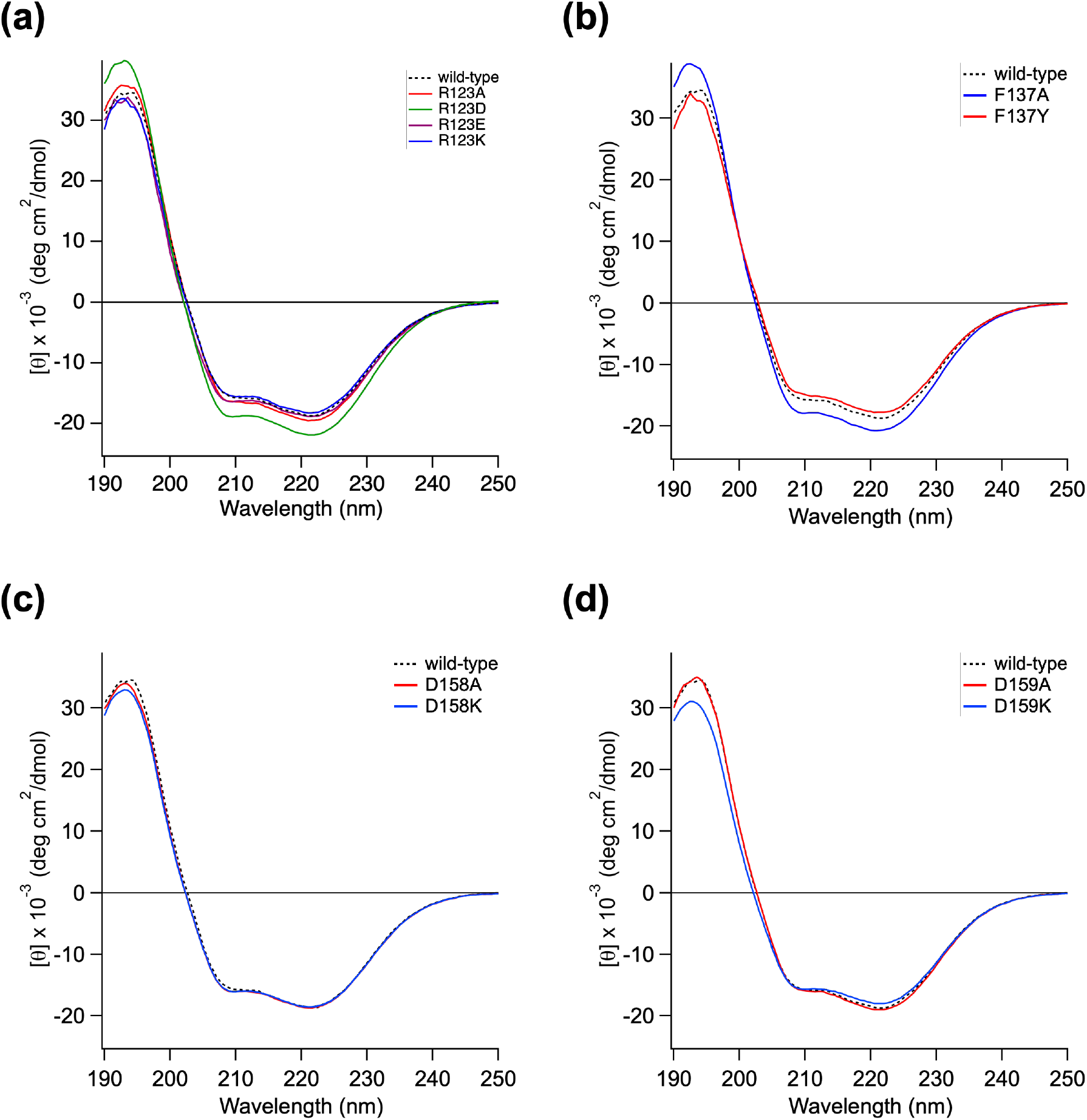
CD spectra of Adk mutants in 5 mM potassium phosphate buffer (pH 8.0) at 25°C; (a) R123X mutants; (b) F137X mutants; (c) D158X mutants; (d) D159X mutants. The spectra shown is the averages of two measurements.

**Table 2.**
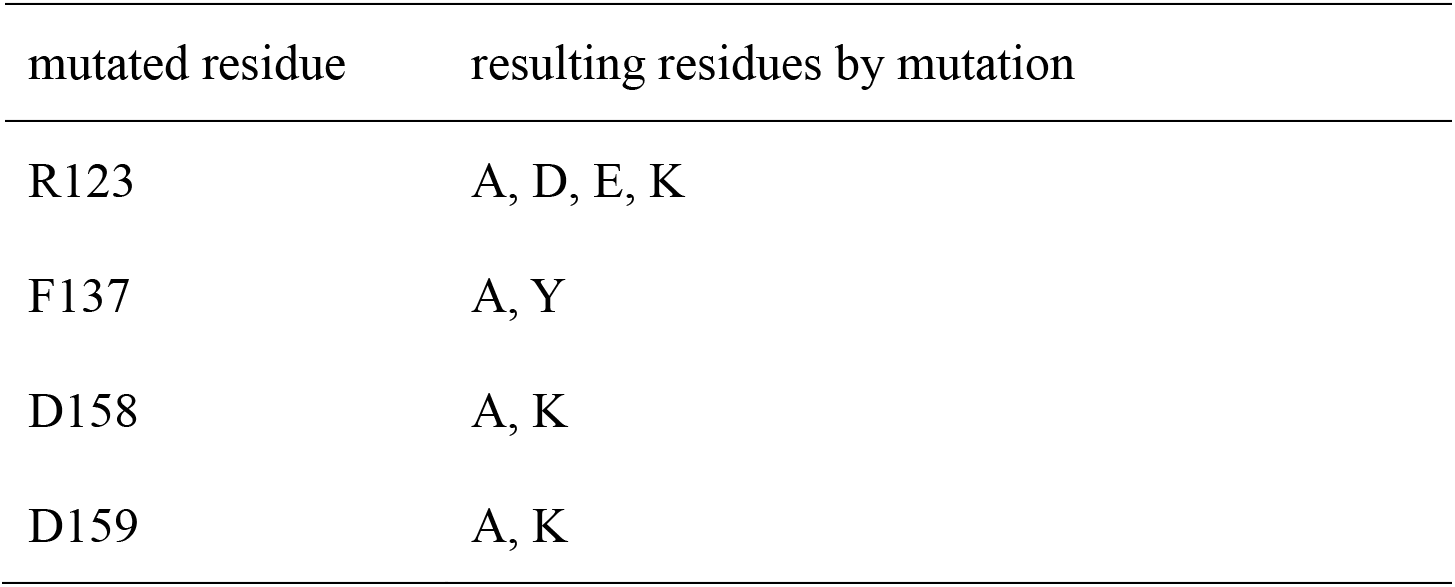
Mutations for Adk.

### Binding affinities of wtAdk and its mutants to OP-4

To evaluate the affinities of the Adk mutants for OP-4, we performed isothermal titration calorimetry (ITC) measurements. Representative thermograms and titration curves are shown in Fig. 5. The data for mutants other than presented in Fig. 5 are demonstrated in Figs. S1–S4. Dissociation constants (*K*_d_) and thermodynamic parameters obtained from the ITC titration curves are listed in Table 3.

**Fig. 5.**
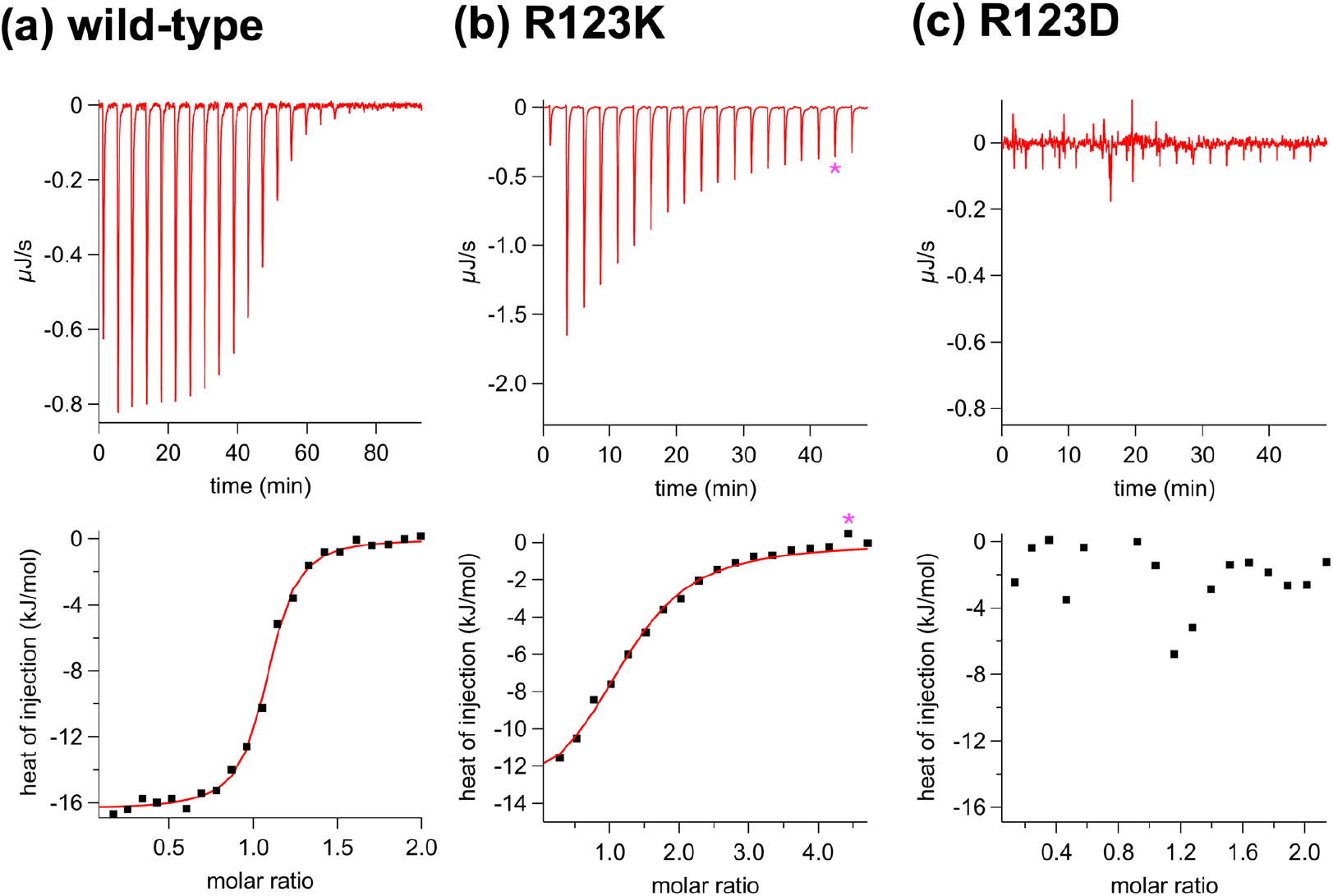
Representative isothermal titration calorimetry (ITC) thermograms and titration curves; (a) wtAdk; (b) R123K mutant; (c) R123D mutant. Measurement conditions: 20 mM Tris-HCl (pH 8.0) containing 150 mM NaCl at 25°C. Fitting lines were drawn based on 1:1-binding model. In figure (b), the peak marked with an asterisk (*) was not used for curve fitting due to large deviations greater than 3σ.

**Table 3.** ITC analysis for Adk.

| Adk | $N^b$ | $K_d^c$<br>(nM) | $\Delta G$<br>(kJ/mol) | $\Delta H$<br>(kJ/mol) | $T\Delta S^d$<br>(kJ/mol) |
| --- | --- | --- | --- | --- | --- |
| wild-type | $1.05 \pm 0.01$ | $73 \pm 8$ | $-40.8 \pm 0.3$ | $-16.4 \pm 0.1$ | $24.4 \pm 0.4$ |
| R123A | n.d.<sup>e</sup> | n.d.<sup>e</sup> | n.d.<sup>e</sup> | n.d.<sup>e</sup> | n.d.<sup>e</sup> |
| R123D | n.d.<sup>e</sup> | n.d.<sup>e</sup> | n.d.<sup>e</sup> | n.d.<sup>e</sup> | n.d.<sup>e</sup> |
| R123E | n.d.<sup>e</sup> | n.d.<sup>e</sup> | n.d.<sup>e</sup> | n.d.<sup>e</sup> | n.d.<sup>e</sup> |
| R123K | $1.20 \pm 0.06$ | $6600 \pm 1400$ | $-29.6 \pm 0.5$ | $-13.8 \pm 1.0$ | $15.7 \pm 1.6$ |
| F137A | $0.88 \pm 0.09$ | $6800 \pm 2400$ | $-29.5 \pm 0.9$ | $-8.5 \pm 1.2$ | $21.1 \pm 2.1$ |
| F137Y | $1.15 \pm 0.01$ | $183 \pm 22$ | $-38.5 \pm 0.3$ | $-17.1 \pm 0.2$ | $21.4 \pm 0.5$ |
| D158A | $1.11 \pm 0.02$ | $290 \pm 62$ | $-37.3 \pm 0.5$ | $-28.5 \pm 0.7$ | $8.9 \pm 1.2$ |
| D158K | $1.05 \pm 0.02$ | $780 \pm 95$ | $-34.9 \pm 0.3$ | $-49.4 \pm 1.0$ | $-14.5 \pm 1.3$ |
| D159A | $1.05 \pm 0.04$ | $1620 \pm 350$ | $-33.1 \pm 0.5$ | $-26.7 \pm 1.7$ | $6.3 \pm 2.2$ |
| D159K | $1.15 \pm 0.08$ | $34000 \pm 3000$ | $-25.5 \pm 0.2$ | $-52.9 \pm 4.7$ | $-27.3 \pm 4.9$ |

<sup>a</sup>20 mM Tris-HCl (pH 8.0) at 25°C. <sup>b</sup>number of binding site. <sup>c</sup>dissociation constant ( $1/K_d = \exp(-\Delta G/RT)$ ). <sup>d</sup>calculated from experimentally determined $\Delta G$ and $\Delta H$ in titration curves obtained at $T = 298.15$ . <sup>e</sup>not determined because no distinct titration curves were observed.

As shown in Fig. 5(a), wtAdk displayed a sigmoidal titration curve analyzable by the 1:1-binding model (*i.e*., *N* ∼1). R123K, F137X, D158X, and D159X mutants also followed the 1:1-binding model (Figs. 5(b) and S1–S3). In contrast, R123A, R123D, and R123E mutants (mutants with a neutral or acidic sidechain) did not display distinct heat changes during titration (Figs. 5(c) and S4), preventing us from analyzing the isothermal titration. Accordingly, we investigated the binding of these mutants to OP-4 using size exclusion chromatography (SEC) and compared the chromogram patterns with that observed for wtAdk (Fig. S5). In wtAdk, the protein with OP-4 was detected as a fraction with a smaller elution volume than free wtAdk (Fig. S5(a)), whereas no such SEC elution shift was observed for the R123 mutants.

The binding affinity of wtAdk against OP-4 was determined to be *K*_d_ = 73 ± 8 nM with a negative enthalpy and positive entropy changes. The mutants that display distinct isothermal titration curves also showed exothermic complexation. R123K, F137A, D159A, and D159K mutants showed 20–500-fold increases in *K*_d_-values. In contrast, the increases in the *K*_d_-values of two D158 mutants were moderate compared to those of the former mutants. For F137Y, the increase in the *K*_d_-value was limited to less than threefold. Positive entropic changes were observed in wtAdk and most mutants, whereas D158K and D159K showed negative entropic changes.

### MD simulations for Adk

To discuss the effects of mutation modes on the affinities and thermodynamic parameters in OP-4 binding, we performed MD simulation to estimate the solution structures of wt- and mutant Adk prior to OP-4 binding (*i.e*., initial states of the binding processes). We focused on R123 and D159 for MD simulations because these mutants showed larger reduction of OP-4-binding affinity than other mutants.

The crystal structure of wtAdk (Fig. 6(a)) indicates the sidechains of R123 and D159 are separated by 3.0 Å. R156 is located near D159 with distances of *ca.* 3.8 Å. Because R156, D158, and D159 are on a flexible loop, their side chains are expected to interact dynamically in solution. Accordingly, we focused on these residues to perform trajectory analyses based on the MD simulations.

**Fig. 6.**
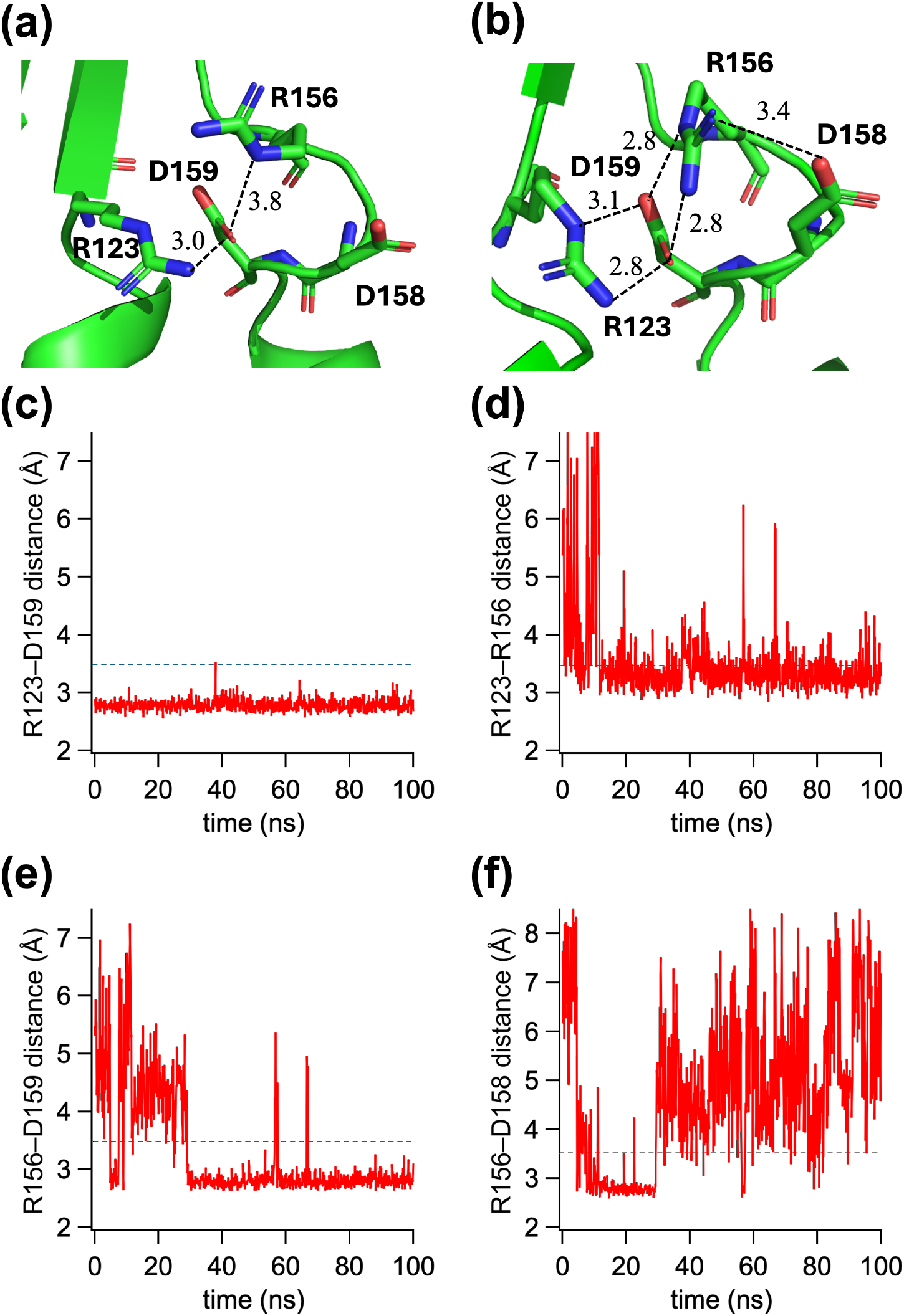
Molecular dynamics (MD) simulation for free wtAdk; (a) crystal structure of wtAdk (initial structure of MD simulation; PDB: 4AKE); (b) a representative snapshot in MD simulation (taken at 50 ns); (c) trajectory of inter-side-chain distance between R123 and D159; (d) trajectory of inter-side-chain distance between R123 and R156; (c) trajectory of inter-side-chain distance between R156 and D159; (d) trajectory of inter-side-chain distance between R156 and D158; In figures (a) and (b), minimum heavy-atom distances between side chains are indicated in angstrom (Å) unit. In figures (c)–(f), dashed lines indicate the 3.5 Å cutoff used for occupancy (OCC) calculations.

Time-dependent changes in inter-sidechain distance (ISD) and occupancy (OCC) were analyzed, where OCC was defined as the probability that two sidechains remain within 3.5 Å during the simulation. A representative snapshot structure during the MD simulation for wtAdk is shown in Fig. 6(b). The trajectories of inter-side-chain distances in wtAdk are presented in Figs. 6(c)–6(f). These analyses were also performed for R123, D158, and D159 mutants in the same manner as done for wtAdk. Average ISDs and OCCs values are summarized in Table 4. Trajectories ISDs for the mutants are shown in Figs. S6–S10. The changes in Cα-RMSDs (root-mean-square deviations) during MD calculations for all the Adk variants are shown in Fig. S11.

**Table 4.** Trajectory analysis of MD simulations for Adk.

| Adk | average of ISD<sup>b</sup> and OCC<sup>c</sup> |  |  |  |
| --- | --- | --- | --- | --- |
|  | residue 123 /<br>residue 159 | residue 123 /<br>residue 156 | residue 156 /<br>residue 159 | residue 156 /<br>residue 158 |
| wild-type | 2.79 Å<br>(OCC >99%) | 3.51 Å<br>(OCC 71%) | 3.27 Å<br>(OCC 75%) | 4.80 Å<br>(OCC 24%) |
| R123K | 6.89 Å<br>(OCC 4.4%) | 6.54 Å<br>(OCC 1.0%) | 2.84 Å<br>(OCC 98%) | 5.76 Å<br>(OCC 5.6%) |
| D158A | 2.82 Å<br>(OCC 99%) | 4.32 Å<br>(OCC 45%) | 3.70 Å<br>(OCC 72%) | 5.26 Å<br>(OCC 5.0%) |
| D158K | 2.82 Å<br>(OCC >99%) | 5.46 Å<br>(OCC 53%) | 4.57 Å<br>(OCC 34%) | 7.49 Å<br>(OCC 0.1%) |
| D159A | 7.86 Å<br>(OCC 0.19%) | 6.78 Å<br>(OCC 3.1%) | 6.73 Å<br>(OCC 0.57%) | 3.99 Å<br>(OCC 65%) |
| D159K | 8.16 Å<br>(OCC 0.09%) | 11.1 Å<br>(OCC 0.09%) | 5.67 Å<br>(OCC 5.4%) | 3.55 Å<br>(OCC 76%) |

<sup>a</sup>MD calculations were performed using AMBER14 force field, an aqueous solution model with 0.9% NaCl aq ion concentration, point charges assigned at pH = 8.0 at 298.15 K, and additional Na<sup>+</sup> or Cl<sup>-</sup>. The trajectory was sampled every 250 ps. <sup>b</sup>For residue pairs, the guanidinium nitrogen atoms (NE, NH1, and NH2) in an arginine residue, the carboxylate oxygen atoms (OD1 and OD2) in an aspartic acid residue, the amine nitrogen atom (NZ) in a lysine residue and the methyl carbon in alanine were used in calculation. <sup>c</sup>obtained by counting the number of frames that satisfies the distance criterion ( $\leq 3.5$ Å) and dividing by the total number of analyzed frames.

A representative snapshot during the MD simulation for wtAdk displayed interactions between R123 and D159 with distances of 2.8–3.1 Å (Fig. 6(b)), together with interactions between D159 and R156. The average ISD between R123 and D159 is 2.79 Å with an OCC value exceeding 99%. The average ISDs of R123–R156 and R156–D159 pairs are within the distance criterion (≤3.5 Å). In contrast, the average ISD for the R156–D158 pair was 4.80 Å with an OCC value of 24%.

In the R123K mutant, K123–D159 pair has an ISD of 6.98 Å with an OCC of 4.4%; namely, the mutation of R123 to Lys (K) decreased the interaction possibility. For K123–R156 and R156–D158 pairs, the average ISDs of these pairs are also larger than 3.5 Å with OCCs of less than 10%. In contrast, the ISD of K156–D159 is 2.84 Å, which is smaller than that observed in wtAdk.

In D158A and D158K mutants, their average ISDs and OCCs for R123–D159 pair (average ISD: 2.82 Å; OCC: >99%) are similar to those for wtAdk. In contrast, their average ISDs for the R123–R156 and R156–D159 pairs increased from that of wtAdk to larger values than the distance criterion (≤3.5 Å). The OCCs values for these pairs also decreased from those of wtAdk. For R156–A158 pair in D158A mutant, the OCC value dropped down to 5.0%. Furthermore, R156–K158 pair in D158K mutant showed an average ISD of 7.49 Å with an OCC of 0.1%, due to the electrostatic repulsion between the two basic residues.

The mutations of D159 caused the increases in average ISDs and the decreases in OCCs for the pairs involved by residue 159 (*i.e.*, residues 123–159 and 156–159 pairs). In contrast, the R156–D158 pairs in D159 mutants were found to be the smaller average ISDs with larger OCCs, compared to those observed in wtAdk. The trajectories of ISD for R156–D158 pair (Figs. S9(d) and S10(d)) also indicate that the distance between these residues tend to be within the distance criterion (≤3.5 Å) during the simulation time.

### Comparison of crystal structure with the binding model constructed by SAXS data

The overlapped figures of the crystal structure with DENSS algorithm-calculated electron density map derived from the previously acquired SEC-SAXS data [14] is shown in Fig. S12. Excess density was observed in regions not fully accounted for by the crystal structure.

## Discussion

### Crystal structure of Adk:OP-4 complex and OP-4-binding modes

The crystal structure revealed that OP-4 binds to the surface formed by the expanded LID and CORE domains in Adk. This binding mode is consistent with our previous epitope-mapping study: the chemical modification at residues 55 (NMP domain) and 169 (near the boundary of CORE and LID domains) does not interfere with the OP-4 binding [14]. The crystal structure showed that OP- 4 binds to the site apart from these residues.

R123 in Adk, a residue involved in the complex formation, is known to be one of the first contact residues for ATP-binding [32]. However, its primary role appears to be associated with the conformational transition rather than ATP-binding itself. A previous study reported that the mutation of R123 into alanine decreases the *k*_cat_-value to 1% of that observed in wtAdk, although the *K*_m_-value for ATP (obtained under saturating AMP concentrations) is less affected by the mutation [33]; the ATP-binding ability is retained even the mutation at R123. Consistent with these findings, our previous ^31^P-NMR and titration experiments demonstrated that ATP can still bind to Adk in the presence of OP-4 [15], indicating that the OP-4-binding site is not completely overlapped with the ATP-binding site. On the OP-4 side, the interacting residues are located within the diversified regions in the “side-and-loop library”. This indicates that the engineered binding surface directly participates in target recognition. The OP-4-binding site shown in the crystal structure differs from that in the complex model proposed in our previous SEC-SAXS study. This issue will be discussed later.

### Availability of mutational approach

Similarity in CD spectra between wtAdk and Adk mutants (Fig. 4) assures the availability of mutational approach for elucidating the contribution of above-mentioned residues to the complexation, without considering influences on the protein secondary structure by mutations.

### Outline of mutation effects on OP-4 binding to Adk

The negative enthalpic change observed for wtAdk (Fig. 5(a), Table 2) reflects a salt-bridge formation and several hydrogen-bonding interactions as shown in the crystal structure. A positive entropic change can be attributed to desolvation on the complex formation [34]. Complex formations involving entropic advantages have also been shown in the nucleotide-binding to wtAdk [35] and in other examples of antibody mimetic-involving protein-protein interactions [36–38].

The mutations at residues that are related with the complex formation caused the reduction of OP-4 binding (*i.e*., the increased *K*_d_-values) or non-observation of distinct heat changes. This finding indicates that the interactions presented in the crystal structure were abolished by the mutations. In other words, the interactions occurred at the mutated residues occur also in solution. The changes in complexation profiles (*i.e.*, *K*_d_-values and thermodynamic parameters) depended on mutation modes. This matter on each residue will be discussed in detail below.

Although the crystal structure of free wtAdk (Fig. 6(a)) showed no distinct salt-bridge between the sidechains of R123 and D159, a snapshot taken in the MD simulation (Fig. 6(b)) displayed a hydrogen bond network among R123–D159–R156 residues with ISDs of 2.8–3.1 Å. The trajectory of ISD for R123–D159 pair (Fig. 6(c)) also demonstrated a persistent ion pair during the simulation time around 2.8 Å. Furthermore, the trajectories of R123–R156 and R156–D159 pairs (Figs. 6(d) and (e)) also indicated that these residues tend to frequently interact with each other in solution, with distances of less than 3.5 Å. Therefore, wtAdk forms the “R123–D159–R156 hydrogen bond network” in solution.

The mutation of R123 *or* D159 cause more than 20-fold increases in *K*_d_-values. Some R123 mutants prevented us to correctly determine *K*_d_-values because of no distinct titration curves. Therefore, the structure with the “R123–D159–R156 hydrogen bond network” is essential to the binding of OP-4 and the mutations at these residues may cause critical decreases in the binding ability.

### Effects of R123 in Adk on the complex formation

R123 participates in multiple interactions between Adk and OP-4 in the crystal structure. On the mutation of R123 to alanine, aspartic acid or glutamic acid, neither a distinct heat change in ITC measurements nor elution shift in SEC study was observed. This result indicates that OP-4 hardly binds to these Adk mutants because a salt-bridge interaction with D75 of OP-4 is abolished. The salt-bridge interaction between the guanidium of R123 with D75 in OP-4 is the most critical factor to achieve the complex formation.

Although both arginine and lysine have a positively charged sidechains, the R123K mutant displayed a 90-fold larger *K*_d_-value, compared with wtAdk. The increase in *K*_d_-value is mainly caused by the loss of entropic advantage (*TΔS* = 24.4 kJ/mol for wtAdk; 15.7 kJ/mol for the R123K mutant). The MD simulation for the R123K mutant suggested little possibility of K123–D159 interaction (ISD = 6.89 Å; OCC: 4.4%; Fig. S6(a)). In turn, D159 tends to interact with R156 (OCC: 98%). This structural configuration requires a rearrangement of sidechain orientations in K123/D159/R156 residues to form a suitable conformation for the OP-4 binding. Such a structural rearrangement decreases entropic gain in the complex formation.

### Effects of F137 in Adk on complex formation

F137 in Adk contacts with Y31 in OP-4 through π– π stacking in the crystal structure. An 86-fold increase in *K*_d_-value on the replacement of F137 with alanine is mainly caused by an enthalpic penalty, attributable to the loss of π– π interactions. The increase in *K*_d_-value was moderate in the F137Y mutant (less than 3-fold increase) because an aromatic sidechain is retained even after mutation to tyrosine; the π– π stacking interaction is still possible before and after the mutation.

### Effects of D158 in Adk on complex formation

Changes in *K*_d_-value by mutations at D158 were within a 10-fold increases, which is moderate compared to the changes observed in the R123 mutations. The increases in *K*_d_-value observed for D158 mutants are mainly caused by the loss of entropic advantage, indicating that D158 mutations led to structural rearrangements for the complex formation. MD simulations for D158 mutants showed the persistent R123– D159 interaction (ISDs = 2.82 Å with an OCC of >99%). However, R123–R156 and R156– D159 interactions tend to be weakened, featured by larger ISDs than the distance criterion. The decreases of the interactions between these pairs extended the interaction strength between R156 and residue 158. These results suggest that D158 indirectly regulates the orientation of the R156 sidechain to assist the retainment of “R123–D159–R156 hydrogen bond network”. A negative entropy change observed in the D158K mutant is due to electrostatic repulsion between R156 and K158, requiring a large conformational change to pre-organize the structure before OP-4 binding.

### Effects of D159 in Adk on complex formation

Since D159 forms an ion pair with R123, the increase in *K*_d_-value by mutations at D159 can be explained by the disruption of the R123– D159–R156 hydrogen-bond network. The MD simulations for D159 mutants also indicate little possibility of interactions between R123, R156, and residue 159 (Figs. S10(a), (b), and (c)). In turn, R156 tends to interact with D158, featured by the increased OCCs of R156–D158 pair from that of wtAdk. The reduction of binding affinity on D159 mutations is due to the entropic disadvantage. Especially, the negatively large entropy in D159K mutant is due to the local clustering of three basic residues (*i.e.*, R123, R156, and K159), where the orientation of R123 is different from that observed in wtAdk to relieve electrostatic repulsion. Therefore, a large conformational rearrangement is required for the D159K Adk mutant to form the complex with OP-4.

### Difference in OP-4-binding modes between in crystal structure and structural model proposed by SEC-SAXS data

Comparison of the crystal structure with the DENSS- calculated electron density map derived from the previous SEC-SAXS data revealed excess electron density regions that were not fully accounted for by the crystal structure (Fig. S12). One possible explanation for this discrepancy is conformational heterogeneity and/or dynamic properties of the Adk:OP-4 complex in solution. Nevertheless, the mutational, ITC, and MD analyses consistently support the interaction modes observed in the crystal structure. These results suggest that OP-4 binds to the interface identified crystallographically in solution as well as in the crystal lattice.

## Conclusion

We successfully determined the crystal structure of Adk:monobody OP-4 complex with the aid of hierarchical clustering analysis. The structure revealed the OP-4 binding to the surface formed by the expanded LID and CORE domains in Adk. This binding mode is consistent with our previous epitope-mapping study, although it differs from the OP-4-binding site proposed in the previous SEC-SAXS study. Mutational, ITC, and MD analyses consistently support the crystallographically observed OP-4-binding mode. Particularly, R123 was identified as a key residue for OP-4 binding, as several R123 mutants almost lost their ability in the complex formation with OP-4. The MD simulations further suggested that the R123–D159–R156 hydrogen-bond network is largely retained in solution, which may facilitate efficient OP-4 binding.

The discrepancy between the OP-4-binding sites suggested by the crystal structure and the SAXS-derived model remains unresolved. Future high-resolution SEC-SAXS analyses, including measurements using synchrotron radiation, may provide additional insights into the solution-state architecture of the Adk–OP-4 complex.

## Acknowledgements

This work was financially supported by Grant-in-Aid for Scientific Research (C) (Japan Society for the Promotion of Science (JSPS) KAKENHI grants JP22K05316 to TM, JP21K05386 to ST, and JP24K08717 to ST), a Grant-in-Aid for Transformative Research Area (A) (JSPS KAKENHI grant JP23H04559 to ST), a Grant-in-Aid for Early-Career Scientists (JSPS KAKENHI grant JP25K18416 to HM), Murata memorial foundation to TM, Izumi Science and Technology Foundation to TM, and Iketani Science and Technology Foundation to IN and TM. This work was also supported by JST SPRING, Japan (Grant Number JPMJSP2140 to IN). The authors are grateful to the staff at beamline BL45XU, SPring-8, Japan (Proposal No. 2023B2523). The authors thank Prof. Shun Hirota (NAIST) for his kind arrangement of our facility usage.

## Author contributions

NO, KY, KO performed all experiments described in this paper and wrote the manuscript. IN prepared the crystal of Adk:OP-4 complex and wrote the manuscript. STF and HM performed hierarchical clustering analysis for X-ray diffraction data and wrote the manuscript. ST resolved the X-ray crystallographic data and wrote the manuscript. ST and TM led this work overall.

**Abbreviations**
Adk: adenylate kinase
Ap_5_A: *P*^1^, *P*^5^-di(adenosine-5’) pentaphosphate
ATP: adenosine triphosphate
CD: circular dichroism
*E. coli*: *Escherichia coli*.
HCA: hierarchical clustering analysis
ITC: isothermal titration calorimetry
NMR: nuclear magnetic resonance
OP-4: a monobody binding to the OPEN-form Adk
RMSD: root mean square deviation
PDB: protein data bank
PPIs: protein-protein interactions
SDS-PAGE: sodium dodecyl sulfate-polyacrylamide gel electrophoresis
SEC: size exclusion chromatography
SEC-SAXS: size exclusion chromatography/small-angle X-ray scattering
TCEP: tris(carboxyethyl)phosphine
TEV: Tobacco Etch Virus
Tris: tris(hydroxymethyl)aminomethane
UV-vis: ultraviolet- visible
wt: wild-type

## Supporting Information

**Table S1.** Amino acid sequences of Adk and monobody OP-4

**Table S2.** Data collection and refinement statistics in X-ray crystallographic analysis of the Adk:OP-4 complex

**Fig. S1.** Isothermal titration calorimetry (ITC) thermograms and titration curves of F137X Adk mutants

**Fig. S2.** Isothermal titration calorimetry (ITC) thermograms and titration curves of D158X Adk mutants

**Fig. S3** Isothermal titration calorimetry (ITC) thermograms and titration curves of D159X Adk mutants

**Fig. S4** Isothermal titration calorimetry (ITC) thermograms and titration curves of R123X Adk mutants

**Fig. S5** Size exclusion chromatograms of wtAdk and R123X mutants with or without OP-4

**Fig. S6** Trajectories of inter-side-chain distances in MD simulation for Adk R123K mutant

**Fig. S7** Trajectories of inter-side-chain distances in MD simulation for Adk D158A mutant

**Fig. S8** Trajectories of inter-side-chain distances in MD simulation for Adk D158K mutant

**Fig. S9** Trajectories of inter-side-chain distances in MD simulation for Adk D159A mutant

**Fig. S10** Trajectories of inter-side-chain distances in MD simulation for Adk D159K mutant

**Fig. S11** Trajectory of Cα-RMSD (root-mean-square deviations) during MD simulation

**Fig. S12** Overlapping of the crystal structure and DENSS-calculated electron density map for the Adk:OP-4 complex

