## Supplemental Tables S1, S2, Figs. S1-S12 for "Structural basis for monobody OP-4 binding to open-form adenylate kinase"

**Contents**

| | | |
|---|---|---|
| **Table S1** | Amino acid sequences of Adk and monobody OP-4 | 3 |
| **Table S2** | Data collection and refinement statistics in X-ray crystallographic analysis of the Adk:OP-4 complex | 4 |
| **Fig. S1** | Isothermal titration calorimetry (ITC) thermograms and titration curves of F137X Adk mutants | 5 |
| **Fig. S2** | Isothermal titration calorimetry (ITC) thermograms and titration curves of D158X Adk mutants | 6 |
| **Fig. S3** | Isothermal titration calorimetry (ITC) thermograms and titration curves of D159X Adk mutants | 7 |
| **Fig. S4** | Isothermal titration calorimetry (ITC) thermograms and titration curves of R123X Adk mutants | 8 |
| **Fig. S5** | Size exclusion chromatograms of wtAdk and R123X mutants with or without OP-4 | 9 |
| **Fig. S6** | Trajectories of inter-side-chain distances in MD simulation for Adk R123K mutant | 10 |

| | | |
|---|---|---|
| **Fig. S7** | Trajectories of inter-side-chain distances in MD simulation for Adk D158A mutant | 11 |
| **Fig. S8** | Trajectories of inter-side-chain distances in MD simulation for Adk D158K mutant | 12 |
| **Fig. S9** | Trajectories of inter-side-chain distances in MD simulation for Adk D159A mutant | 13 |
| **Fig. S10** | Trajectories of inter-side-chain distances in MD simulation for Adk D159K mutant | 14 |
| **Fig. S11** | Trajectory of Cα-RMSD (root-mean-square deviations) during MD simulation | 15 |
| **Fig. S12** | Overlapping of the crystal structure and DENSS-calculated electron density map for the Adk:OP-4 complex | 16 |

**Table S1.** Amino acid sequences of Adk and monobody OP-4

| protein | amino acid sequence |
| --- | --- |
| Adk[a] | MRIILLGAPG AGKGTQAQFI MEKYGIPQIS TGDMLRAAVK SGSELGKQAK DIMDAGKLVT DELVIALVKE RIAQEDCRNG FLLDGFPRTI PQADAMKEAG INVDYVLEFD VPDELIVDRI VGRRVHAPSG RVYHVKFNPP KVEGKDDVTG EELTTRKDDQ EETVRKRLVE YHQMTAPLIG YYSKEAEAGN TKYAKVDGTK PVAEVRADLE KILG |
| OP-4[b] | (GS) VSSVPTKLEV VAATPTSLLI SWDAPAVTVY YYIITYGETG RNASFQEFAV PGSKSTATIS GLSPGVDYTI TVYADGCGYD LCSPISINYR T |

[a]Amino acid residues marked in red were mutated in this study. [b]*N*-Terminus-attached "GS" part (marked by blankets "( )") is a fragment derived from the TEV protease-cleavage in purification. Colored parts are residues diversified in phage display selections (red: *C* strand; blue: *CD* loop; green: *D* strand; purple: *FG* loop).

**Table S2.** Data collection and refinement statistics in X-ray crystallographic analysis of the Adk:OP-4 complex.

| **Data set** | **Adk : OP-4 complex** |
|---|---|
| X-ray Source | SPring-8 BL45XU |
| wavelength (Å) | 1.0000 |
| temperature (K) | 100 |
| space group | C2 |
| unit-cell parameters | |
| $a$, $b$, $c$ (Å) | 109.51, 42.76, 66.36 |
| $\alpha$, $\beta$, $\gamma$ (°) | 90, 95.86, 90 |
| resolution range (Å) | 50.00–2.53 (2.68–2.53) |
| total reflections | 33180 (5351) |
| unique reflections | 9965 (1566) |
| completeness (%) | 95.5 (95.7) |
| $R_{\mathrm{meas}}$ (%) | 16.5 (93.5) |
| $R_{\mathrm{merge}}$ (%) $^{b}$ | 14.1 (79.4) |
| $CC_{1/2}$ | 0.988 (0.547) |
| solvent content (%) | 47.6 |
| **Refinement** | |
| resolution range (Å) | 44.29–2.53 (2.68–2.53) |
| reflections (working set) | 9963 (975) |
| reflections (test set) | 997 (98) |
| $R_{\mathrm{work}}$ | 0.219 (0.304) |
| $R_{\mathrm{free}}$ | 0.284 (0.379) |
| No. atoms | |
| protein | 2320 |
| water | 0 |
| other | 0 |
| R.m.s. deviation $^{c}$ | |
| Bond length (Å) | 0.009 |
| Bond angles (°) | 1.146 |
| Average $B$ factors (Å$^2$) | |
| protein | 61.52 / 62.91 $^{d}$ |
| water | – |
| other | – |
| Ramachandran (%) | |
| Favored | 95.99 |
| Allowed | 4.01 |
| Disallowed | 0 |

[a]Values in parentheses are for the highest resolution shells.
[b]$R_{\mathrm{merge}}(I) = \Sigma|I(k) - \langle I\rangle| / \Sigma I(k)$, where $I(k)$ is the value of the $k$th measurement of the intensity of a reflection, $\langle I\rangle$ denotes the mean intensity derived from all observations of a specific reflection and its symmetry-related partners. [c]root-mean-square deviation. [d]Average $B$-factors of the Adk / monobody OP-4.

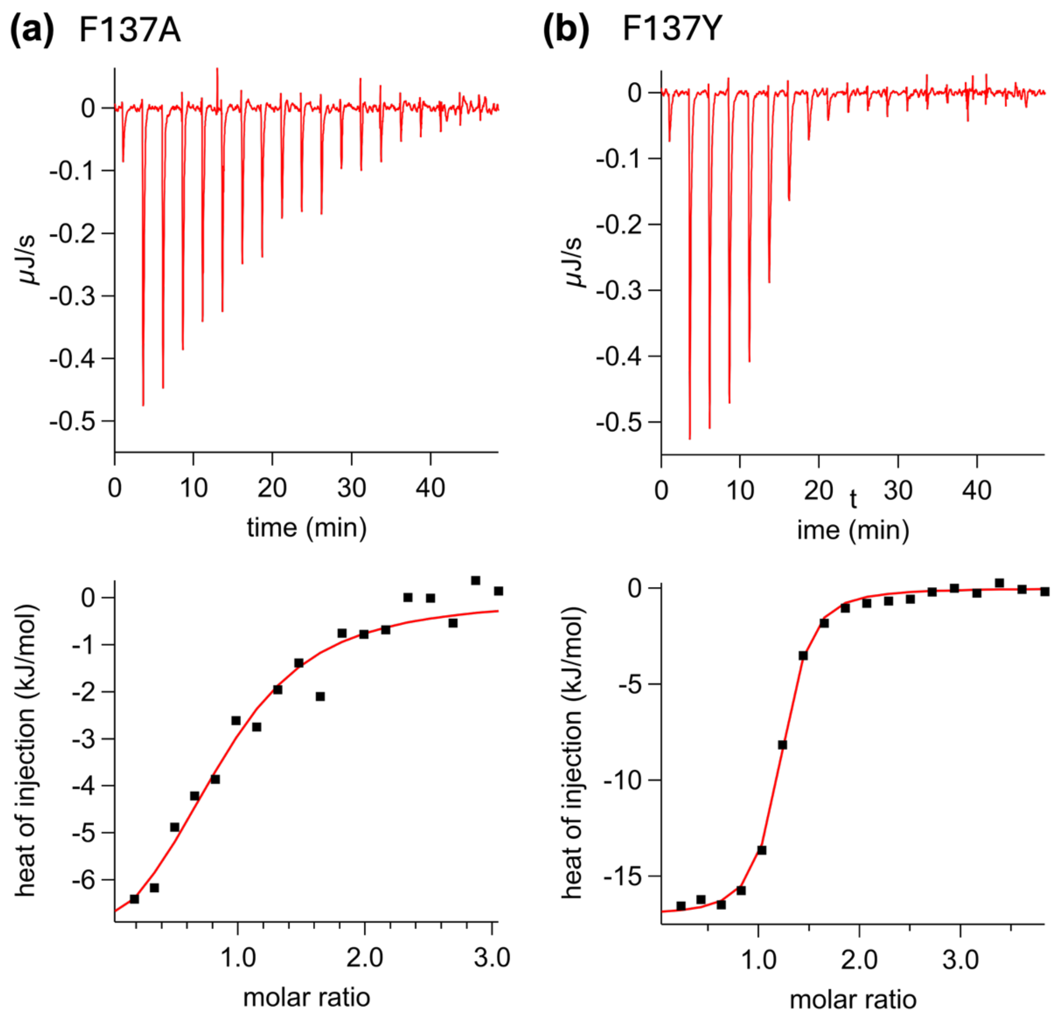


**Fig. S1.** Isothermal titration calorimetry (ITC) thermograms and titration curves of F137X Adk mutants; (a) F137A mutant; (b) F137Y mutant. Fitting lines were drawn based on 1:1-binding model. Measurement conditions: 20 mM Tris-HCl (pH 8.0) containing 150 mM NaCl at 25°C.

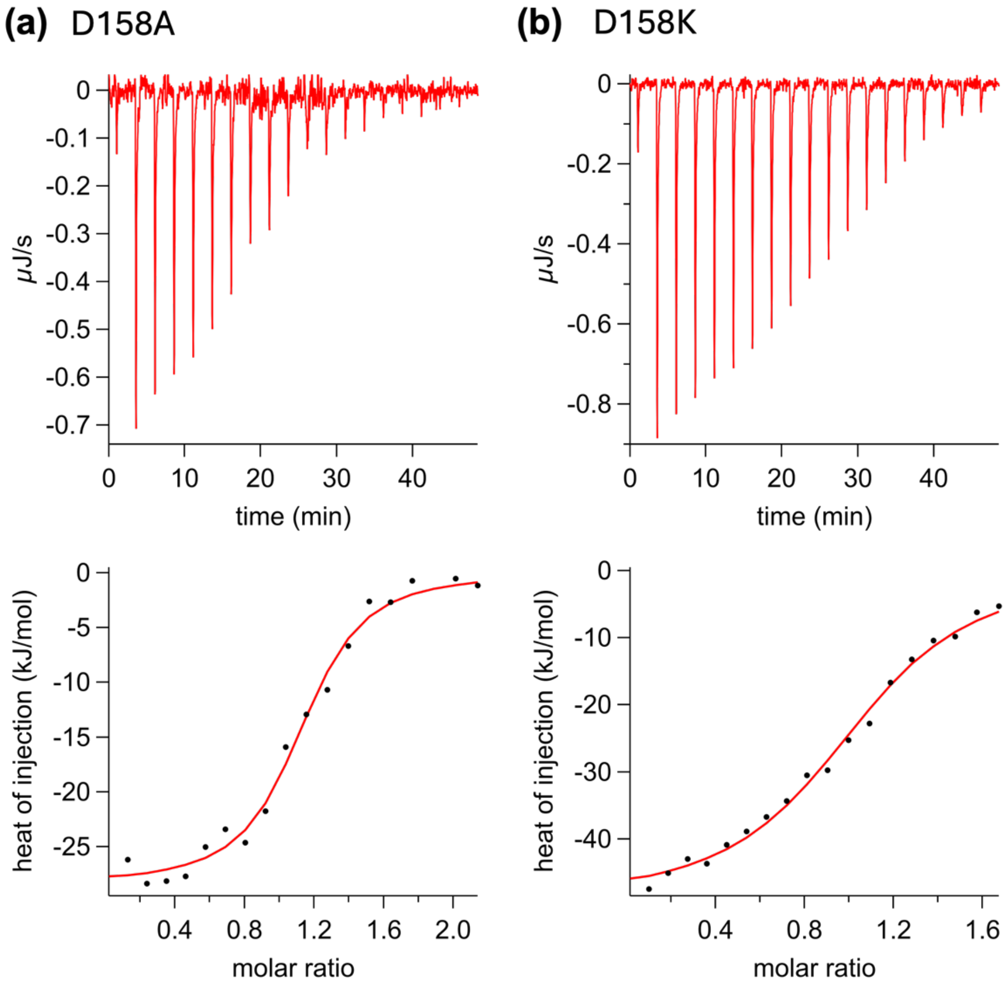


**Fig. S2.** Isothermal titration calorimetry (ITC) thermograms and titration curves of D158X Adk muants; (a) D158A mutant; (b) D158K mutant. Fitting lines were drawn based on 1:1-binding model. Measurement conditions: 20 mM Tris-HCl (pH 8.0) containing 150 mM NaCl at 25°C.

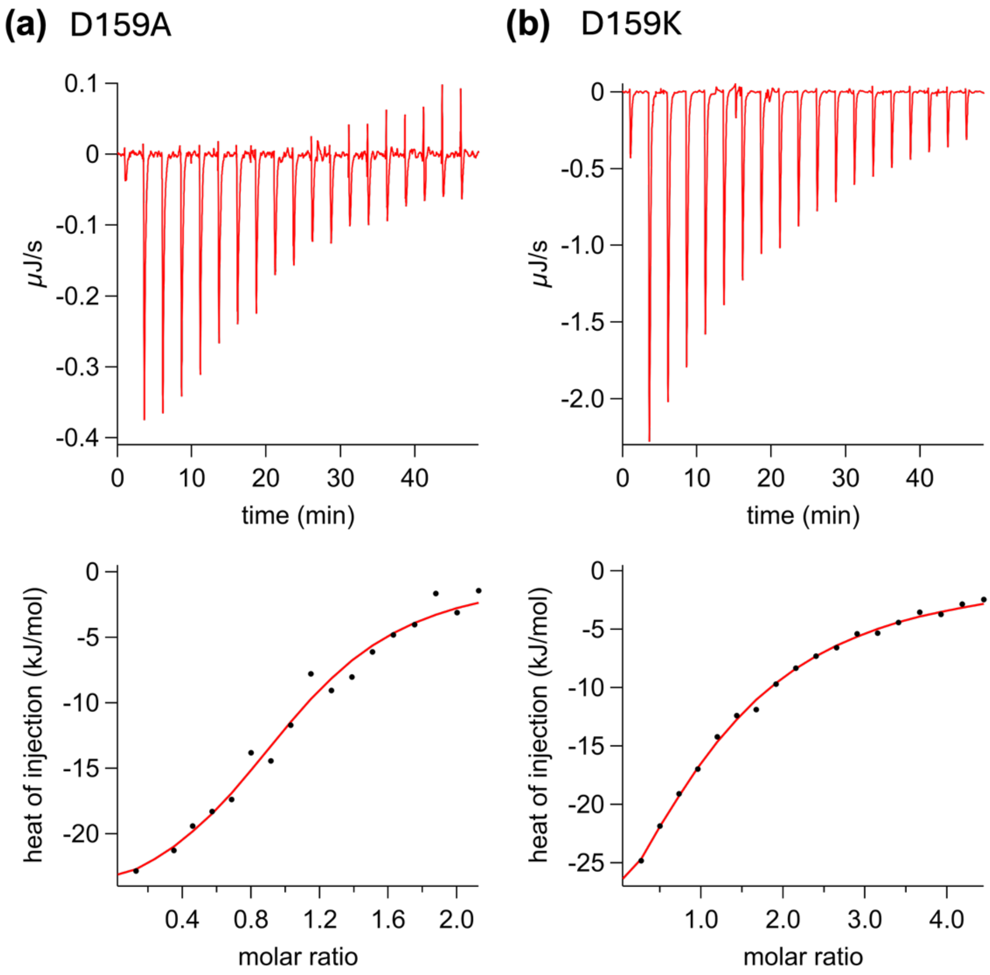


**Fig. S3.** Isothermal titration calorimetry (ITC) thermograms and titration curves of D159X Adk muants; (a) D159A mutant; (b) D159K mutant. Fitting lines were drawn based on 1:1-binding model. Measurement conditions: 20 mM Tris-HCl (pH 8.0) containing 150 mM NaCl at 25°C.

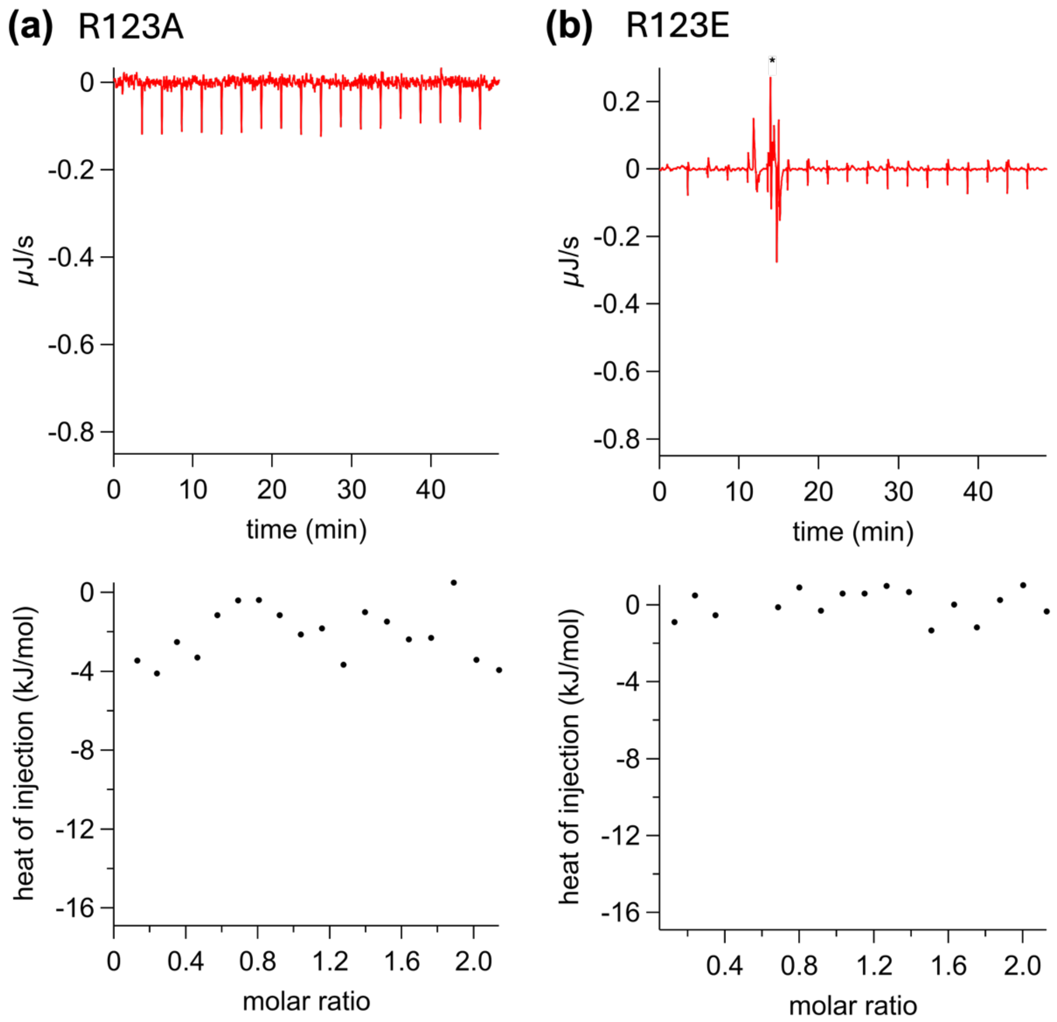


**Fig. S4.** Isothermal titration calorimetry (ITC) thermograms and titration curves of R123X Adk mutants; (a) R123A mutant; (b) R123E mutant. Measurement conditions: 20 mM Tris-HCl (pH 8.0) containing 150 mM NaCl at 25°C.

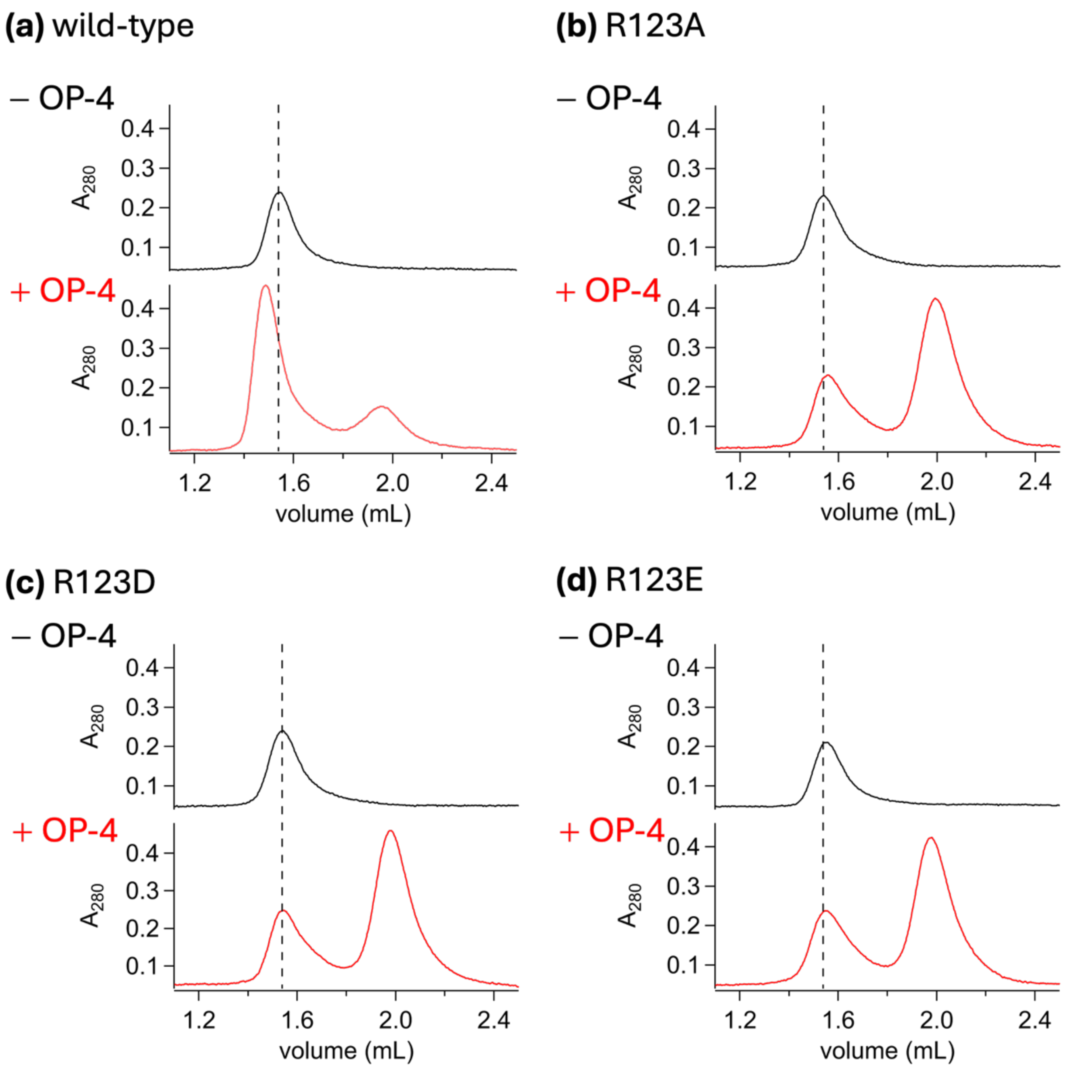


**Fig. S5.** Size exclusion chromatograms of wtAdk and R123X mutants with or without OP-4; (a) wtAdk; (b) R123A mutant; (c) R123D mutant; (d) R123E mutant; Column: Superdex 75 5/150 GL (Cytiva); flow rate : 0.35 mL/min; buffer: 20 mM Tris-HCl (pH 8.0) containing 150 mM NaCl at 25˚C; [Adk] = 30 μM, [OP-4] = 0 or 42 μM. In the presence of OP-4, a peak at large elution volume is assigned as unbound OP-4.

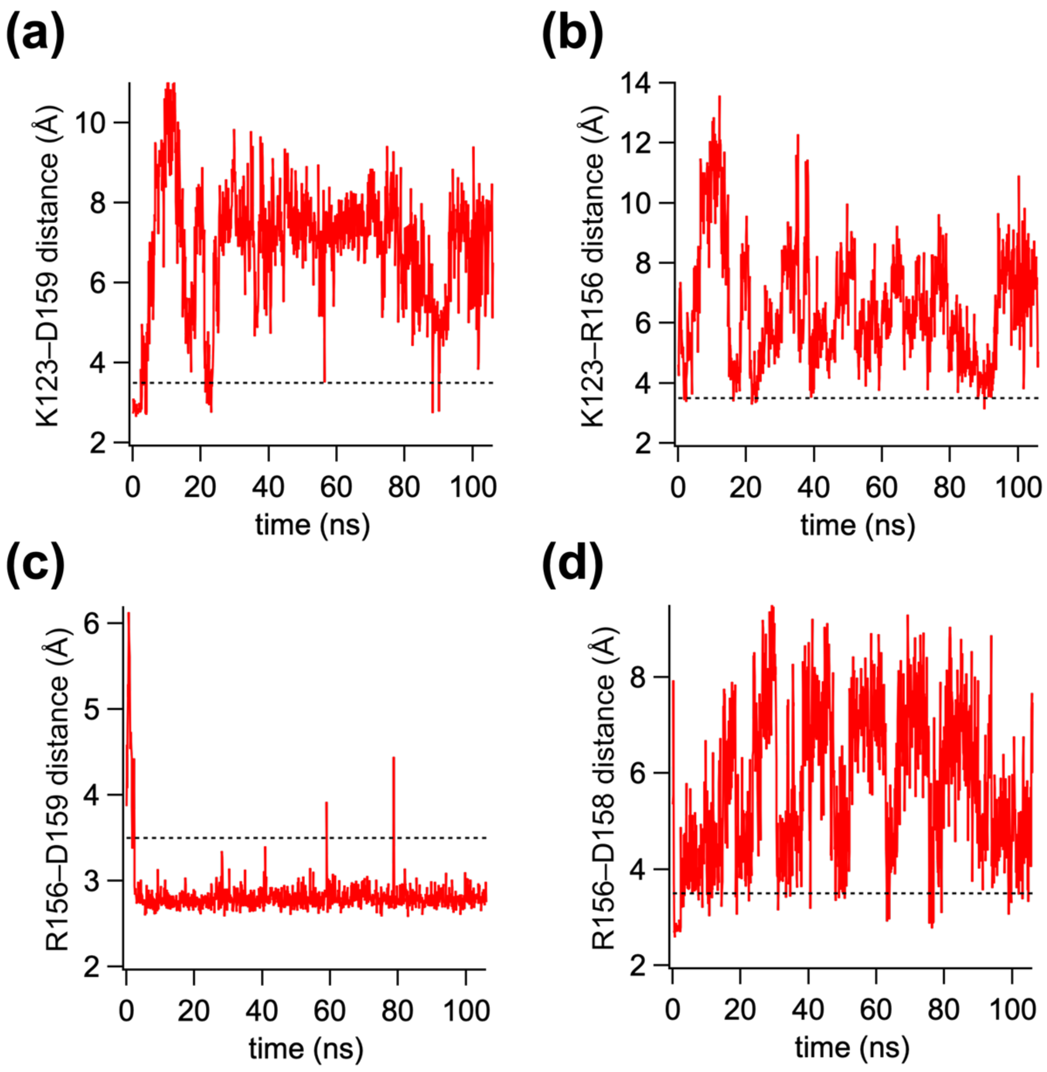


**Fig. S6.** Trajectories of inter-side-chain distances in MD simulation for Adk R123K mutant; (a) K123–D159; (b) K123–R156; (c) R156–D159; (d) R156–D158. Dashed lines indicate the 3.5 Å cutoff used for occupancy (OCC) calculations. Simulation conditions are described in experimental section.

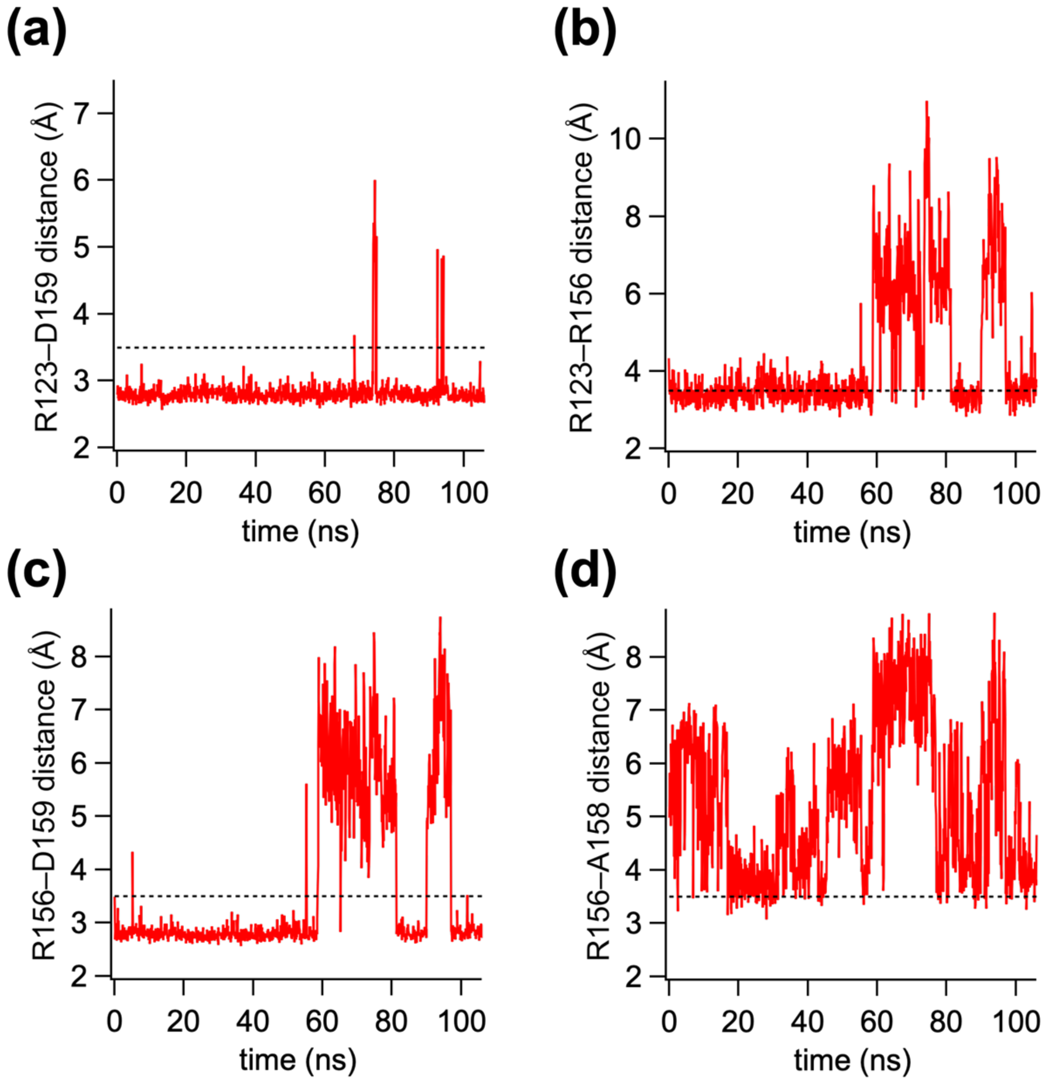


**Fig. S7.** Trajectories of inter-side-chain distances in MD simulation for Adk D158A mutant; (a) R123–D159; (b) R123–R156; (c) R156–D159; (d) R156–A158. Dashed lines indicate the 3.5 Å cutoff used for occupancy (OCC) calculations. Simulation conditions are described in experimental section.

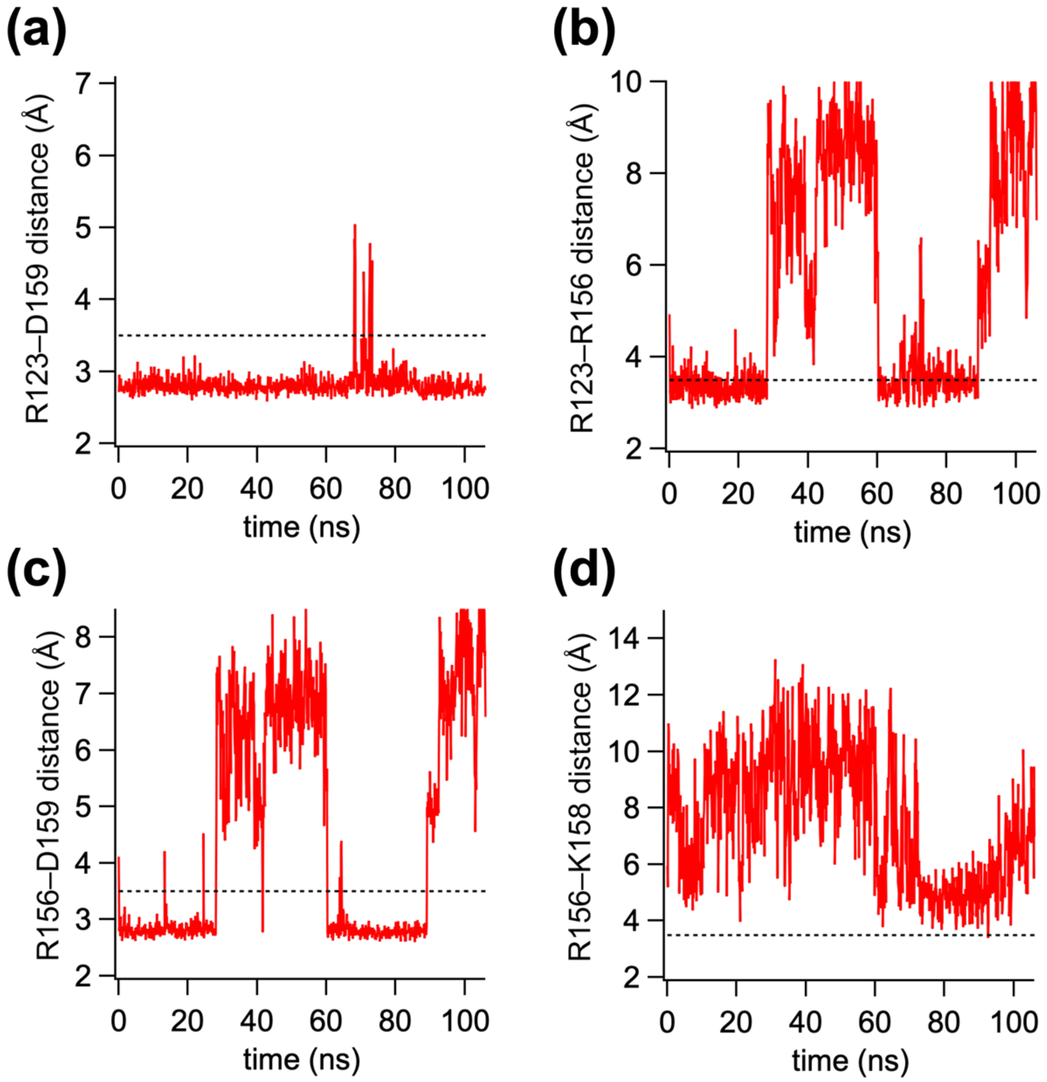


**Fig. S8.** Trajectories of inter-side-chain distances in MD simulation for Adk D158K mutant; (a) R123–D159; (b) R123–R156; (c) R156–D159; (d) R156–K158. Dashed lines indicate the 3.5 Å cutoff used for occupancy (OCC) calculations. Simulation conditions are described in experimental section.

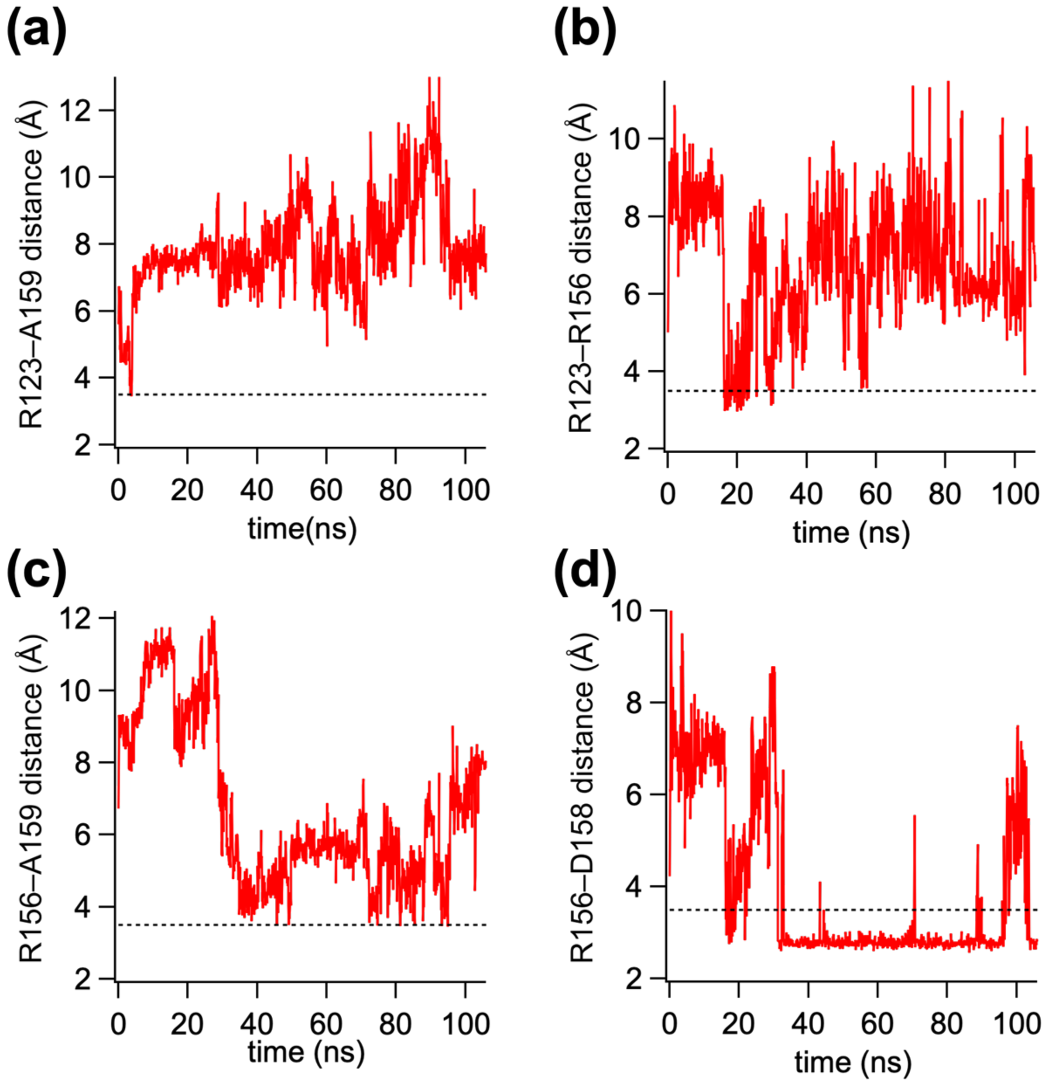


**Fig. S9.** Trajectories of inter-side-chain distances in MD simulation for Adk D159A mutant; (a) R123–A159; (b) R123–R156; (c) R156–A159; (d) R156–D158. Dashed lines indicate the 3.5 Å cutoff used for occupancy (OCC) calculations. Simulation conditions are described in experimental section.

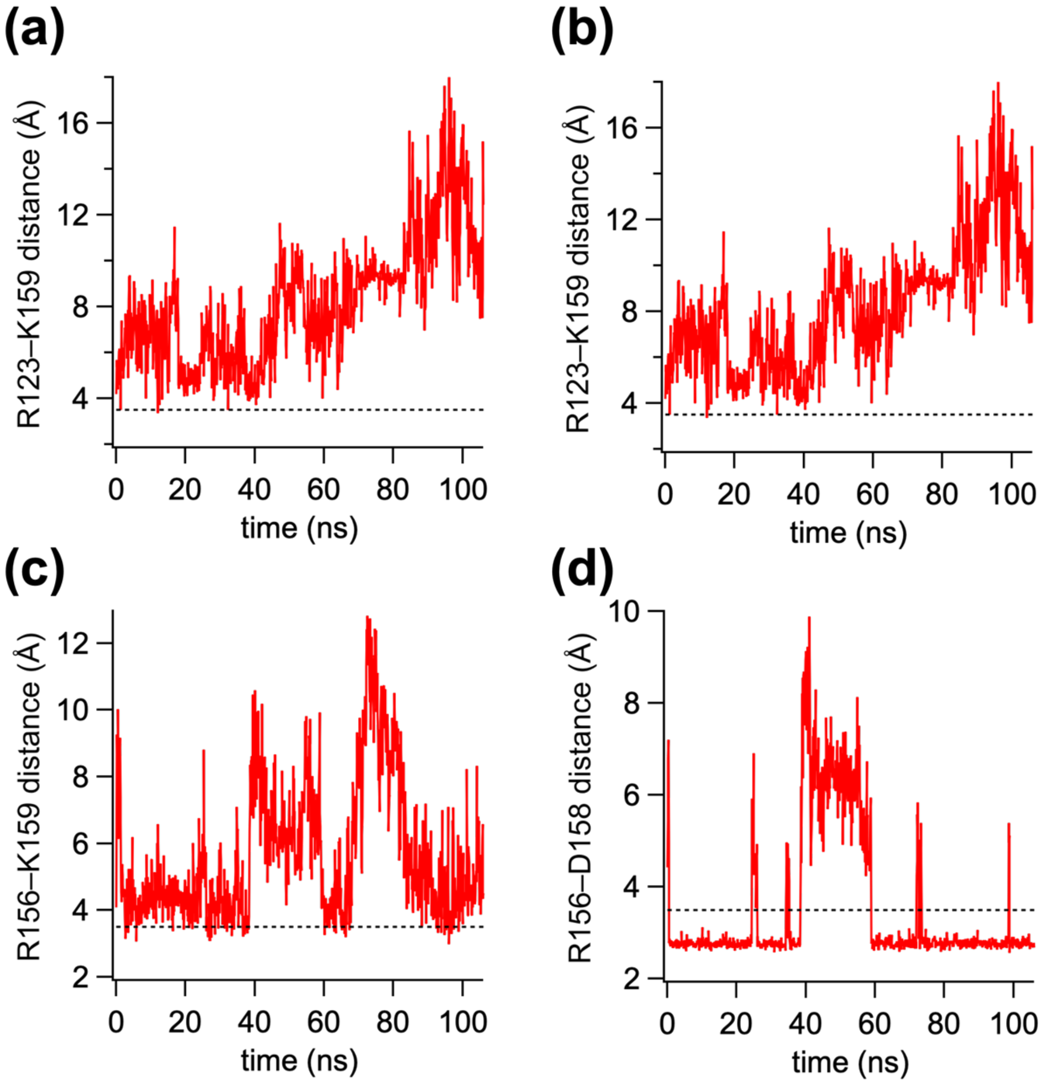


**Fig. S10.** Trajectories of inter-side-chain distances in MD simulation for Adk D159K mutant; (a) R123–K159; (b) R123–R156; (c) R156–K159; (d) R156–D158. Dashed lines indicate the 3.5 Å cutoff used for occupancy (OCC) calculations. Simulation conditions are described in experimental section.

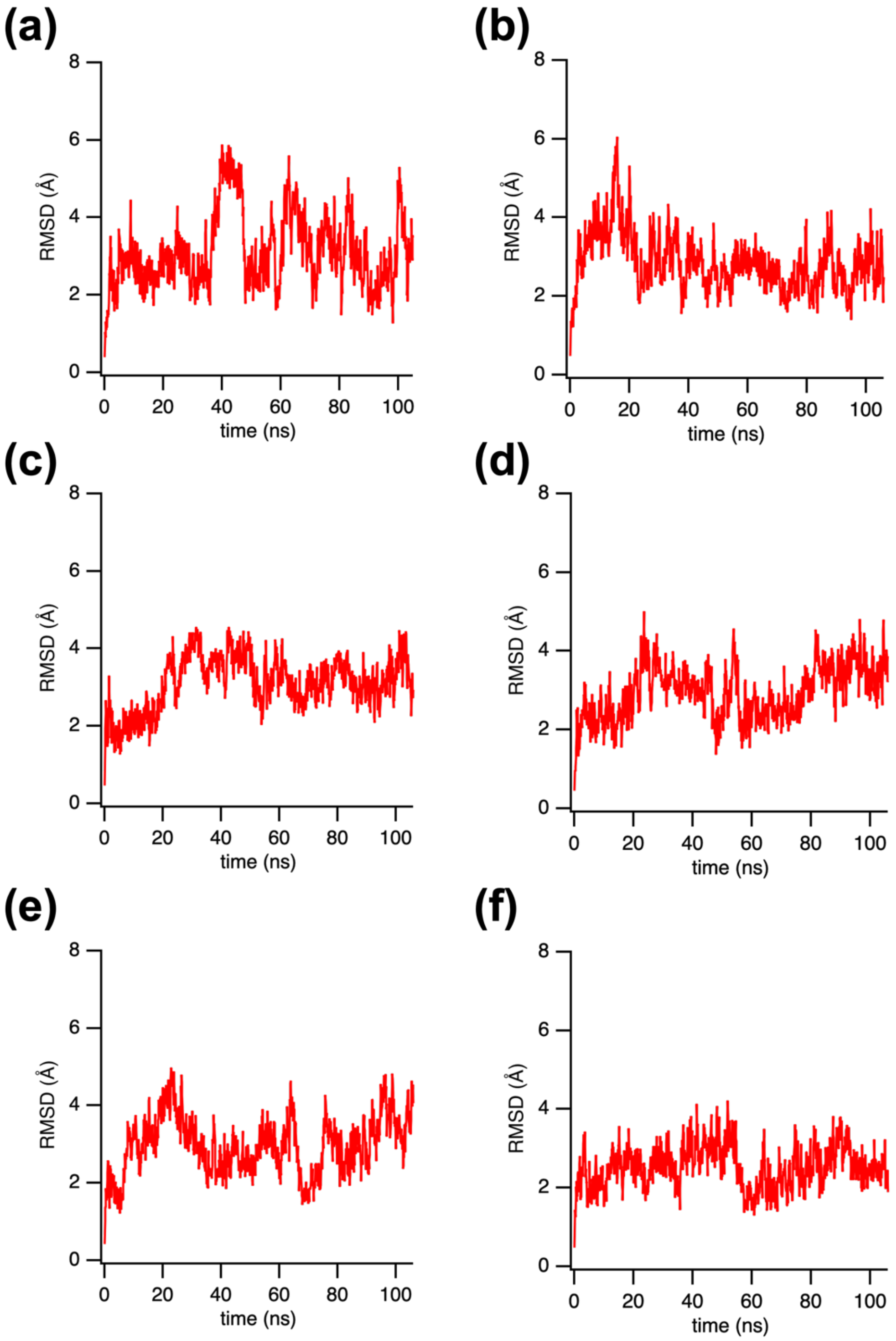


**Fig. S11.** Trajectory of Cα-RMSD (root-mean-square deviations) during MD simulation for the wild-type Adk and its mutants; (a) wild-type; (b) R123K; (c) D158A; (d) D158K; (e) D159A; (f) D159K. Simulation conditions are described in experimental section.

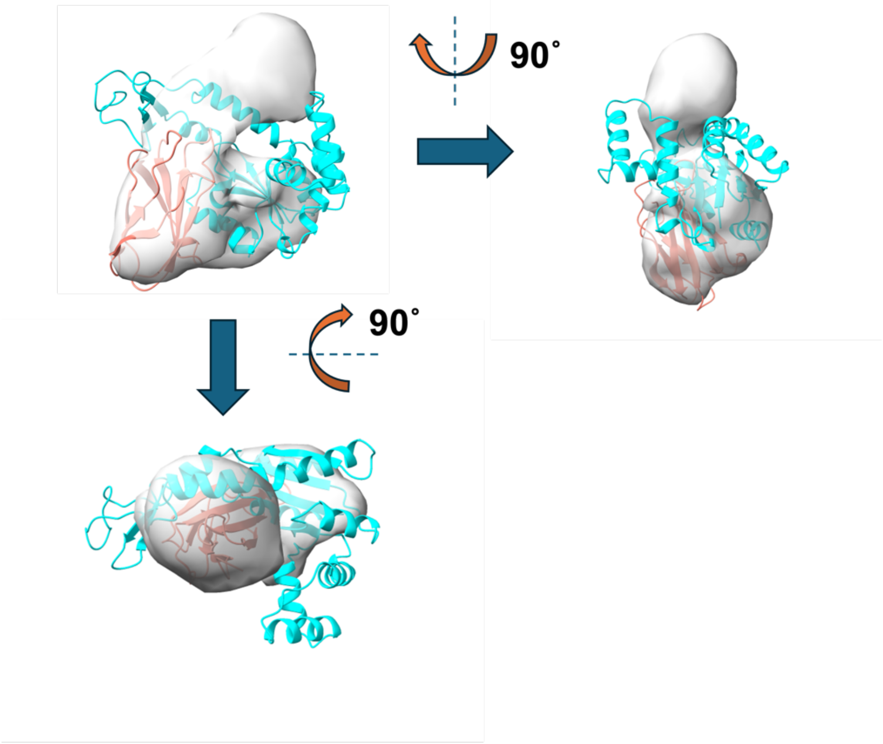


**Fig. S12.** Overlapping of the crystal structure and DENSS-calculated electron density map (0.1 σ) for the Adk:OP-4 complex. The DENSS-calculated electron density map was quoted from the previous work [6]. In the crystal structure, the parts marked in cyan and orange indicate wtAdk and OP-4, respectively.
